# Inhibition of Anti-viral Stress Granule Formation by infectious bronchitis virus endoribonuclease nsp15 Ensures Efficient Virus Replication

**DOI:** 10.1101/2020.06.11.145862

**Authors:** Bo Gao, Xiaoqian Gong, Shouguo Fang, Wenlian Weng, Yingjie Sun, Chunchun Meng, Lei Tan, Cuiping Song, Xusheng Qiu, Weiwei Liu, Maria Forlenza, Chan Ding, Ying Liao

## Abstract

Cytoplasmic stress granules (SGs) are generally triggered by stress-induced translation arrest for storing mRNAs. Recently, it has been shown that SGs exert anti-viral functions due to their involvement in protein synthesis shut off and recruitment of innate immune signaling intermediates. The largest RNA virus, coronavirus, mutates frequently and circulates among animals, imposing great threat to public safety and animal health; however, the significance of SGs in coronavirus infections is largely unknown. Infectious bronchitis virus (IBV) is the first identified coronavirus in 1930s and has been prevalent in poultry farm for many years. In this study, we provide evidence that IBV overcomes the host antiviral response by inhibiting SGs formation *via* the virus-encoded endoribonuclease nsp15. By immunofluorescence analysis, we observed that IBV infection not only did not trigger SGs formation in approximately 80% of the infected cells, but also impaired the formation of SGs triggered by heat shock, sodium arsenite, or NaCl stimuli. We show that the intrinsic endoribonuclease activity of nsp15 is responsible for the inhibition of SGs formation. In fact, nsp15-defective recombinant IBV (rIBV-nsp15-H238A) greatly induced the formation of SGs, along with accumulation of dsRNA and activation of PKR, whereas wild type IBV failed to do so. Consequently, infection with rIBV-nsp15-H238A triggered transcription of IFN-β which in turn greatly affected recombinant virus replication. Further analysis showed that SGs function as antiviral hub, as demonstrated by the attenuated IRF3-IFN response and increased production of IBV in SG-defective cells. Additional evidence includes the aggregation of PRRs and signaling intermediates to the IBV-induced SGs. Collectively, our data demonstrate that the endoribonuclease nsp15 of IBV suppresses the formation of antiviral hub SGs by regulating the accumulation of viral dsRNA and by antagonizing the activation of PKR, eventually ensuring productive virus replication. We speculate that coronaviruses employ similar mechanisms to antagonize the host anti-viral SGs formation for efficient virus replication, as the endoribonuclease function of nsp15 is conserved in all coronaviruses.

**Author summary:** It has been reported that stress granules (SGs) are part of the host cell antiviral response. Not surprisingly, viruses in turn produce an array of antagonists to counteract such host response. Here, we show that IBV inhibits the formation of SGs through its endoribonuclease nsp15, by reducing the accumulation of viral dsRNA, evading the activation of PKR, and by subsequently inhibiting eIF2α phosphorylation and SGs formation. Nsp15 also inhibits SG formation independent of the eIF2α pathway, probably by targeting host mRNA. Depletion of SG scaffold proteins decreases IRF3-IFN response and increases the production of IBV. All coronaviruses encode a conserved endoribonuclease nsp15, and it will be important to determine whether also other (non-avian) coronaviruses limit the formation of anti-viral SGs in a similar manner.

## Introduction

RNA viruses must generate double-stranded RNA (dsRNA) in order to replicate their genome. Host cells consequently employ a variety of pattern recognition (PRRs) receptors to detect dsRNA and trigger innate antiviral responses, which play a pivotal and critical role in fighting viral infections [1]. The host dsRNA-activated protein kinase R (PKR) is a key element of innate antiviral defenses [2]. Following binding to dsRNA, PKR undergoes auto-phosphorylation and phosphorylates the alpha subunit of eukaryotic initiation factor (eIF2α) on serine 51 [2, 3]. Phospho-eIF2α tightly binds to eIF2β, prevents the recycling of ternary complex tRNA^Met^-GTP-eIF2, and inhibits 43S translation complex formation, leading to global translation shut off, severely impairing virus replication [4]. In addition to PKR, there are three other eIF2α kinases involved in translation inhibition: PKR-like endoplasmic reticulum kinase (PERK), general control nonderepressible protein 2 (GCN2), and heme-regulated inhibitor kinase (HRI), which senses unfolded proteins in the endoplasmic reticulum (ER), nutrient starvation/UV[5, 6], and oxidative stress, respectively [7–9]. The translation inhibition leads to polysome disassembly and the subsequent assembly of stress granules (SGs), a membrane-less, highly dynamic warehouse for storing mRNA and translation components [10]. SGs assembly is driven by aggregation-prone cellular RNA-binding proteins: Ras GTPase-activating protein-binding protein 1 (G3BP1), T cell-restricted intracellular antigen 1 (TIA-1), and TIA-1-related protein (TIAR) [11–13]. Meanwhile, the post-translational modifications, including ubiquitination, poly (ADP)-ribosylation, O-linked N-acetyl glucosamination, phosphorylation, and dephosphorylation, regulate the SGs formation by modifying the components of SGs [14, 15]. Once stress is relieved and translation activities are restored, SGs are disassembled and mRNAs rapidly resume translation [16].

In addition to PKR, there are other two groups of PRRs to recognize dsRNA, namely Toll like receptors (TLRs) and RIG-I like receptors (RLRs) [17, 18]. One of the TLRs, TLR3, located on the endosomal membrane, senses dsRNA and single stranded RNA (ssRNA) generated by RNA virus or DNA virus. This in turn activates either the NF-κB or IRF3/7 pathway, resulting in boosting the production of proinflammatory cytokines and type I interferon (IFN) [19]. Another group of essential PRRs, RLRs, composed of RIG-I and MDA5, ubiquitously exist in the cytoplasm of mammal cells, recognize 5’-pppRNA and long dsRNA derived from RNA virus, respectively. Activation of RLRs by viral RNA leads to the aggregation of MAVS and recruitment of a series of signaling intermediates, transmits the signaling to transcription factor IRF3, IRF7, or NF-κB, eventually promoting the transcription of proinflammatory cytokines and type I IFN [20–22]. Consequently, the secretion of type I IFN stimulates the transcription of IFN-stimulated genes (ISGs) *via* the JAK-STAT pathway, which protect neighboring cells from virus infection [23].

Recent evidence has shown that PKR and RLRs are localized to SGs during virus infection [24]. It is proposed that SGs exert specific antiviral activities by providing a platform for interaction between antiviral proteins and non-self RNA. To accomplish efficient replication, some viruses have evolved various mechanisms to circumvent the formation of anti-viral SGs. For instance, Influenza A Virus (IAV) NS1 protein and Vaccinia virus E3L sequester dsRNA from PKR [25, 26], Ebola virus sequesters SG core proteins to viral inclusion body, thereby inhibiting the formation of SGs [27]. For some picornaviruses, leader protease and 3C protease of foot-and-mouth disease virus (FMDV) and 2A protease of Enterovirus 71, disassemble the SGs by cleaving G3BP1 or G3BP2 [28–31]. As for coronaviruses, recent studies show that middle east respiratory syndrome coronavirus (MERS-CoV) 4a accessory protein limits the activation of PKR by binding to dsRNA, thereby inhibiting the formation of SGs and ensuring viral protein translation and efficient virus replication [32, 33]. Mouse hepatitis virus (MHV) replication induces host translational shutoff and mRNA decay, with concomitant formation of stress granules and processing bodies [34]. Porcine transmissible gastroenteritis virus (TGEV) induced SG like granules correlated with viral replication and transcription [35]. Several SG proteins (including caprin and G3BP) have been reported to be associated with IBV-N protein [36]. A recent report shows that infectious bronchitis virus (IBV) infection results in the formation of SGs in approximately 20% of infected cells and inhibits eIF2α-dependent/eIF2α-independent SG formation by unknown mechanisms [37].

Coronaviruses harbor the largest positive-stranded RNA genome among the RNA viruses, with size from 27 kb to 32 kb. The two-third of the 5’ terminus encodes replicase polyproteins (1a and 1ab), while one-third of the 3’ terminus encodes spike protein (S), envelope protein (E), membrane protein (M), nucleocapsid protein (N) and accessory proteins. The proteolysis of overlapped polyproteins is processed by two self-encoded proteases, papain-like protease (PLpro) and 3C-like protease (3CLpro), into 15-16 mature non-structural proteins (nsp1-nsp16). Most of the nsps assemble into a replication and transcription complex (RTC) responsible for virus replication, while several nsps mediate the evasion of host innate immune responses. For example, severe acute respiratory syndrome coronavirus (SARS-CoV) and MERS-CoV nsp1 suppresses host gene expression by mediating host mRNA degradation [38]; the PLpro nsp3 of SARS-CoV and MERS-CoV harbors deubiquitinase activity and interferes with type I IFN responses [39, 40]; feline infectious peritonitis coronavirus (FCoV) and porcine deltacoronavirus (PDCoV) nsp5 inhibits type I IFN response by cleaving NEMO [41, 42]; porcine epidemic diarrhea virus (PEDV) nsp16 restricts IFN production and facilitate virus replication [43]; MHV nsp15 endonuclease activity is key to evade double-stranded RNA (dsRNA) sensing by host sensors and ensures efficient coronavirus replication [44].

IBV is the first identified coronavirus in 1930s and infects avian species [45]. It causes a prevalent disease that has led to substantial economic losses in poultry farm for many decades. Elucidating host responses to IBV infection is fundamental to understand virus replication and identify targets for therapeutic control. In this study, we infected three types of cells with IBV, and found that approximately 80% of infected cells did not display SGs formation. IBV also hindered SGs formation triggered by different canonical stress stimuli. Further analysis showed that IBV nsp15 was involved in the inhibition of SG formation, and that the endoribonuclease activity of nsp15 particularly played a pivotal role. Compared to wild type IBV, infection with the nsp15 endoribonuclease catalytic mutant, rIBV-nsp15-H238A, led to accumulation of higher levels of dsRNA, activation of PKR, and formation SGs, concomitantly with a higher production of IFN-β and lower viral replication. We further demonstrate that SGs play an anti-viral role by using SG-defective cells. To our knowledge, this is the first report describing the role of coronavirus nsp15 in the suppression of integral stress response as well as innate antiviral response.

## Results

### IBV prevents SGs formation in the majority of infected cells

In this study, we employed chicken fibroblast DF-1 cells in most experiments; in specific cases, to facilitate the detection of cellular proteins, and due to the unavailability of antibodies against chicken proteins of interest (see Table 1), we also included mammalian H1299 and Vero cells lines, which also support replication of the IBV-Beaudette strain. To determine whether IBV replication induces SGs formation, H1299, Vero, and DF-1 cells were infected with IBV-Beaudette strain at MOI=1. The occurrence of SGs was assessed by visualizing G3BP1 granules formation while IBV infection was monitored by visualizing the N protein. We determined the kinetics of SGs formation upon infection at 4 hours intervals, by counting the cells with IBV-N protein expression and by calculating the proportion of these that was also positive for G3BP1 granules. In all the three cell types, despite efficient virus infection, as indicated by the expression of N protein and syncytia formation, no SGs formation could be detected from 0 to 8 hours post-infection (h.p.i.), whereas from 12 to 24 h.p.i., G3BP1 granules could be detected, but only in approximately 5%-25% of infected cells (Fig 1A-C). These observations indicate that IBV effectively suppresses SGs formation, and that the inhibition mechanisms employed are not restricted to a specific cell type. In SGs positive cells, another SGs marker, TIAR, was found to colocalize with G3BP1 granules, altogether demonstrating that IBV induces canonical SGs (Fig 1D-E). Due to the lack of antibodies specific for chicken TIAR and lack of cross-reactivity of the anti-human TIAR to chicken TIAR, we displayed the colocalization of TIAR with G3BP1 in Vero and H1299 cells.

**Table 1.** The dilution of primary antibodies and cross reaction to chicken proteins

| name | application | cross-reaction to chicken proteins |
| --- | --- | --- |
| IBV-S | Western blot (1:2000) | - |
| IBV-M | Western blot (1:2000) | - |
| IBV-N | Western blot (1:2000)<br>Immunofluorescence (1:200) | - |
| IBV-nsp3 | Immunofluorescence (1:400) | - |
| G3BP1 | Western blot (1:1000)<br>Immunofluorescence (1:500) | Yes |
| G3BP2 | Western blot (1:1000)<br>Immunofluorescence (1:500) | Yes |
| TIA-1 | Western blot (1:1000) | No |
| TIAR | Immunofluorescence (1:1600) | No |
| PKR | Western blot (1:1000)<br>Immunofluorescence (1:200) | No |
| phospho-PKR | Western blot (1:1000) | No |
| eIF2α | Western blot (1:1000) | No |
| phospho- eIF2α | Western blot (1:1000) | No |
| TBK1 | Western blot (1:1000)<br>Immunofluorescence (1:200) | Yes |
| phospho-TBK1 | Western blot (1:1000) | Yes |
| IRF3 | Western blot (1:1000)<br>Immunofluorescence (1:400) | No |
| phospho-IRF3 | Western blot (1:1000) | No |
| p65 | Western blot (1:1000) | No |
| phospho-p65 | Western blot (1:1000) | No |
| MAVS | Immunofluorescence (1:500) | No |
| MDA5 | Immunofluorescence (1:200) | No |
| TLR3 | Immunofluorescence (1:200) | No |
| TRAF3 | Immunofluorescence (1:200) | No |
| TRAF6 | Immunofluorescence (1:200) | No |
| IKKε | Immunofluorescence (1:200) | No |
| anti-dsRNA J2 | Immunofluorescence (1:100) | - |
| β-actin | Western blot (1:1000) | - |

**Fig 1.**
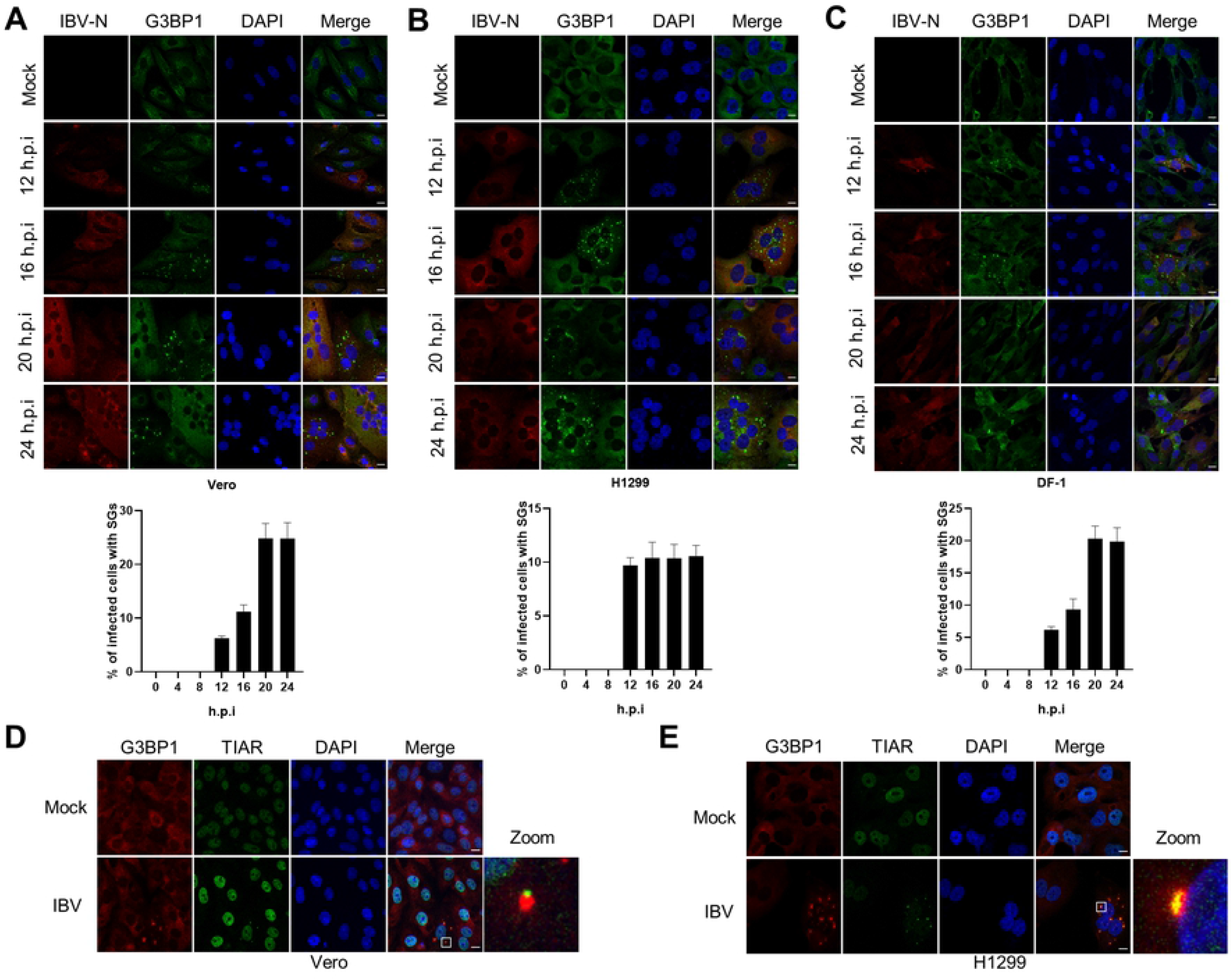
IBV prevents SGs formation in the majority of infected cells. (A-C) Vero, H1299, and DF-1 cells were infected with IBV Beaudette strain at an MOI of 1 or mock infected. At the indicated time points, cells were subjected to immunostaining. Infected cells (red) were identified using a rabbit anti-N protein, SGs (green) with a (cross-reacting) mouse anti-G3BP1 and cell nuclei with DAPI (blue). The SGs positive cells and IBV positive cells in 20 random fields were counted. Bars indicate means ±SD of the percentage of IBV infected cells that were also positive for the presence of SGs. The experiment was performed in triplicate. (D-E) Vero and H1299 cells were infected with IBV as described in A-C. At 20 h.p.i., immunostaining was used to visualize the position of the structural components of SGs using anti-G3BP1 (red) and anti-TIAR (green) antibodies. The enlargement of the inset confirms their co-localization in cytoplasmic granules. Scale bars: 10 μm.

### Phosphorylation status of PKR and eIF2α during IBV infection

Although SGs can be generally induced through eIF2α-dependent or -independent pathways, during viral infections SGs are mainly induced via the PKR-eIF2α pathway. Several past studies demonstrate that some viruses impede SGs formation by preventing the activation of PKR, or by cleaving the SG scaffold protein G3BP1 [46, 47]. To elucidate the inhibition mechanism of SG formation by IBV, we investigated whether IBV interfered with the phosphorylation of PKR and eIF2α or directly affected TIA-1 and G3BP1 protein levels. After having assessed that IBV prevents SGs formation not only in chicken cells but also in two different mammalian cell lines, due to the availability of antibodies directed against mammalian proteins and their lack of cross-reactivity to the chicken proteins of interest, we next proceeded with H1299 and Vero cells. In IBV-infected H1299 cells, PKR and eIF2α phosphorylation was comparable to that observed in mock infected cells (Fig 2A). In Vero cells, however, PKR, but not eIF2α, phosphorylation was slightly increased at 20 and 24 h.p.i. (Fig 2B), altogether suggesting that the PKR-eIF2α-SGs pathway is not obviously triggered by IBV infection. In parallel, we did not observe any cleavage product of either G3BP1 or TIA-1 throughout IBV infection in both H1299 and Vero cells (Fig 2), altogether indicating that IBV may avoid PKR activation to prevent SGs formation.

**Fig 2.**
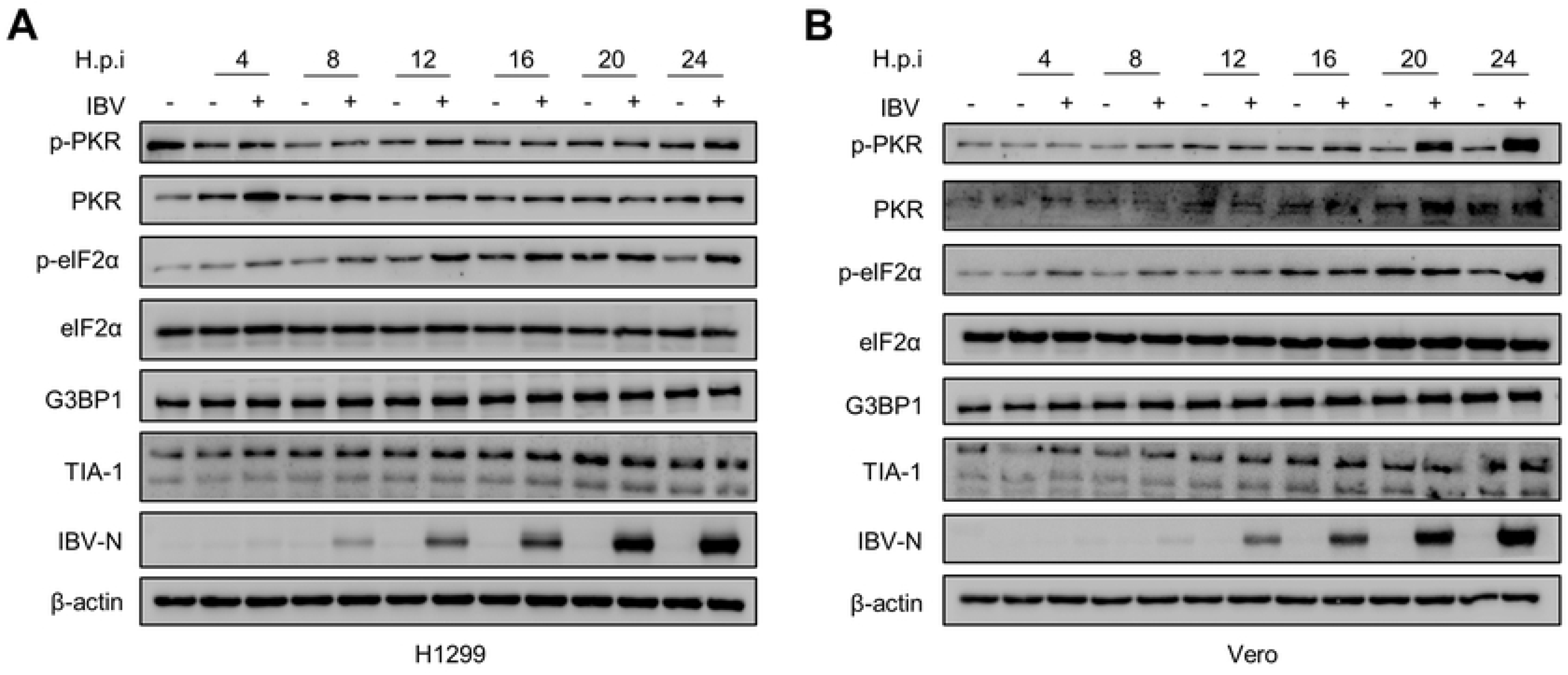
PKR and eIF2α phosphorylation during IBV infection. (A-B) H1299 and Vero cells were infected with IBV at a MOI of 1 or mock infected. Cells were harvested at the indicated time points and processed for western blot analysis using 10 μg total protein per lane. p-PKR, PKR, p-eIF2α, eIF2α, G3BP1, TIA-1, and IBV-N were detected with corresponding antibodies. β-actin was probed as a loading control.

### IBV blocks both eIF2α-dependent and -independent SGs formation

During the course of our study, it was reported that IBV inhibits eIF2α-dependent and -independent SGs induction in Vero cells [37]. Here, we used H1299 and DF-1 cells to further investigate the inhibition of SGs by IBV. H1299 and DF-1 cells were infected with IBV and treated with three different stress stimuli (sodium arsenite, heat shock and NaCl) to induce canonical SGs formation. Sodium arsenite or heat shock promotes phosphorylation of eIF2α in an HRI kinase-dependent manner, leading to translational arrest and subsequent formation of SGs [48], whereas NaCl may enhance the local concentration of mRNAs and cellular proteins by decreasing the cell volume, thereby inducing SGs in an eIF2α-independent manner [49]. In non-infected cells, more than 90% of cells were positive for SGs formation after treatment with these stress stimuli; interestingly, IBV infection prevented SGs formation triggered by these stimuli, as evidenced by the absence of G3BP1 granules exclusively in IBV-positive cells (Fig 3A-C). These results indicate that IBV infection blocks both eIF2α-dependent and independent SGs formation in both, mammalian and avian cells. Sodium arsenite treatment inefficiently triggered G3BP1 aggregation in DF-1 cells due to unknown reasons (data not shown). We next explored whether IBV interfered with the phosphorylation of eIF2α triggered by sodium arsenite or heat shock. We observed a significant upregulation of phospho-eIF2α by sodium arsenite or heat shock treatment in H1299 cells; however, there was no reduction of phospho-eIF2α by IBV infection (Fig 3D-E). Collectively, these data indicate that IBV infection restricts both eIF2α-dependent and -independent SG formation, probably by interfering with SG assembly or disassembly and not with direct eIF2α phosphorylation.

**Fig 3.**
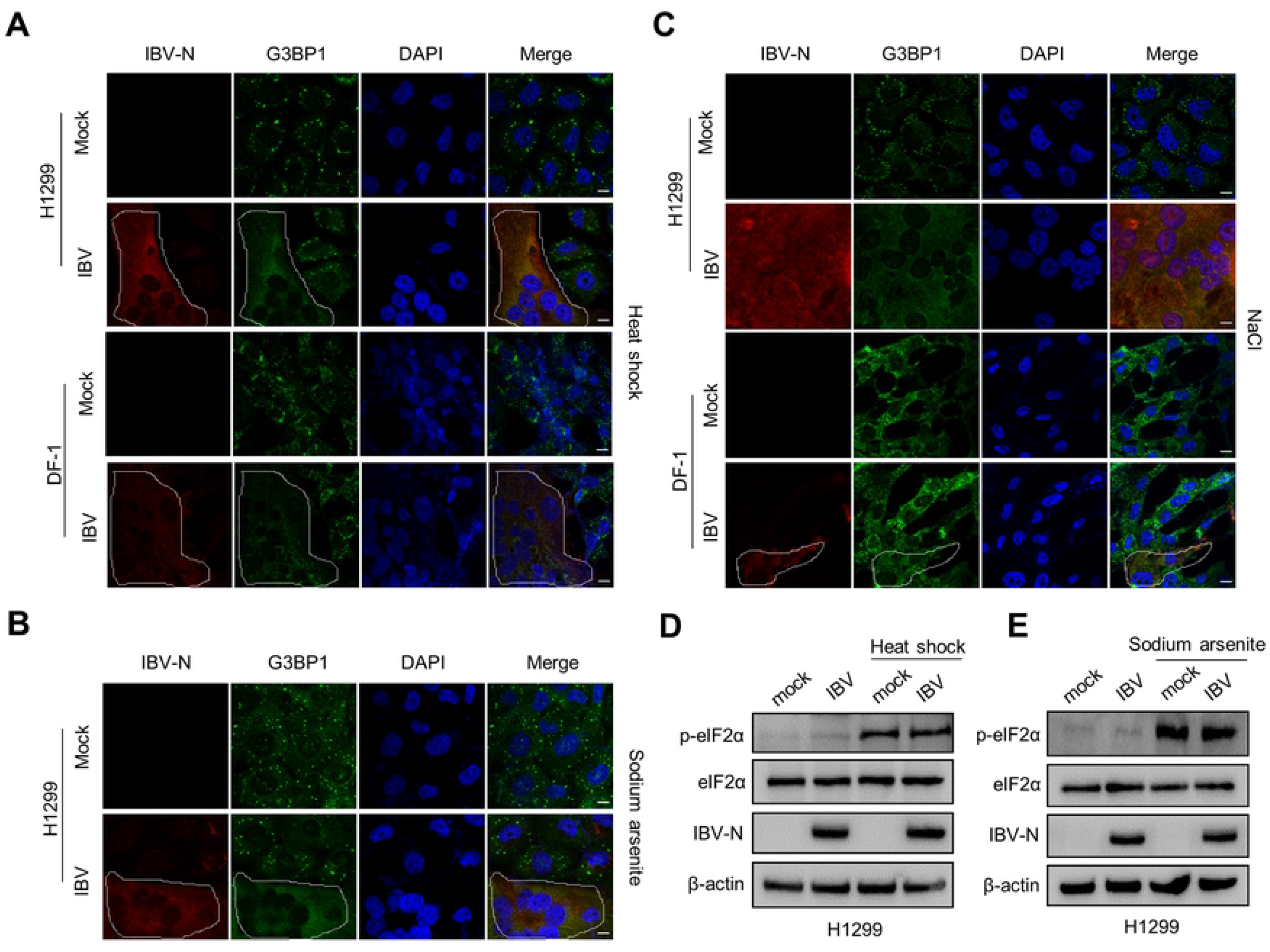
IBV abrogates eIF2α-dependent and -independent formation of SGs. (A-C) H1299 and DF-1 cells were infected with IBV an MOI of 1. At 20 h.p.i., cells received a heat shock treatment (50°C for 20 min) or 1 mM sodium arsenite for 30 min, or 200 mM NaCl for 50 min, followed by immunostaining. Infected cells were detected with anti-N antibody (red), SGs with anti-G3BP1 (green) and cell nuclei with DAPI (blue). Shown are representative images out of three independent experiments. Scale bars: 10 μm. (D-E) H1299 cells were mock infected or infected with IBV and treated with heat shock or sodium arsenite as describe in A and B. At 20 h.p.i., cell lysates (10 μg per lane) were subjected to western blotting analysis to detect p-eIF2α, eIF2α, IBV N, and β-actin.

### Endoribonuclease nsp15 is responsible for inhibition of SG formation

To screen the IBV proteins involved in the suppression of SGs formation, we expressed individual Flag-tagged IBV protein in H1299 cells and triggered the formation of SGs with heat shock. The schematic diagram of proteins encoded by IBV was shown in Fig. 4A. In cells expressing nsp2, nsp4, nsp5, nsp6, nsp7, nsp8, nsp9, nsp12, nsp16, 3b, E, 5a, 5b, M, or N, SGs formation remained intact (Fig 4B), suggesting that alone these proteins have no inhibitory effect on the formation of SGs. Interestingly, only in nsp15-expressing cells, the heat shock-induced SGs were absent (as indicated by white arrow), suggesting that nsp15 may be the viral protein responsible for efficient suppression of SGs formation. We also investigated, but failed to detect, the expression of nsp3, nsp10, nsp13, nsp14, S, and 3a, therefore it cannot be excluded that also these viral proteins might be involved in inhibition of SG formation. These results however demonstrate that nsp15 alone is sufficient to block SGs formation.

**Fig 4.**
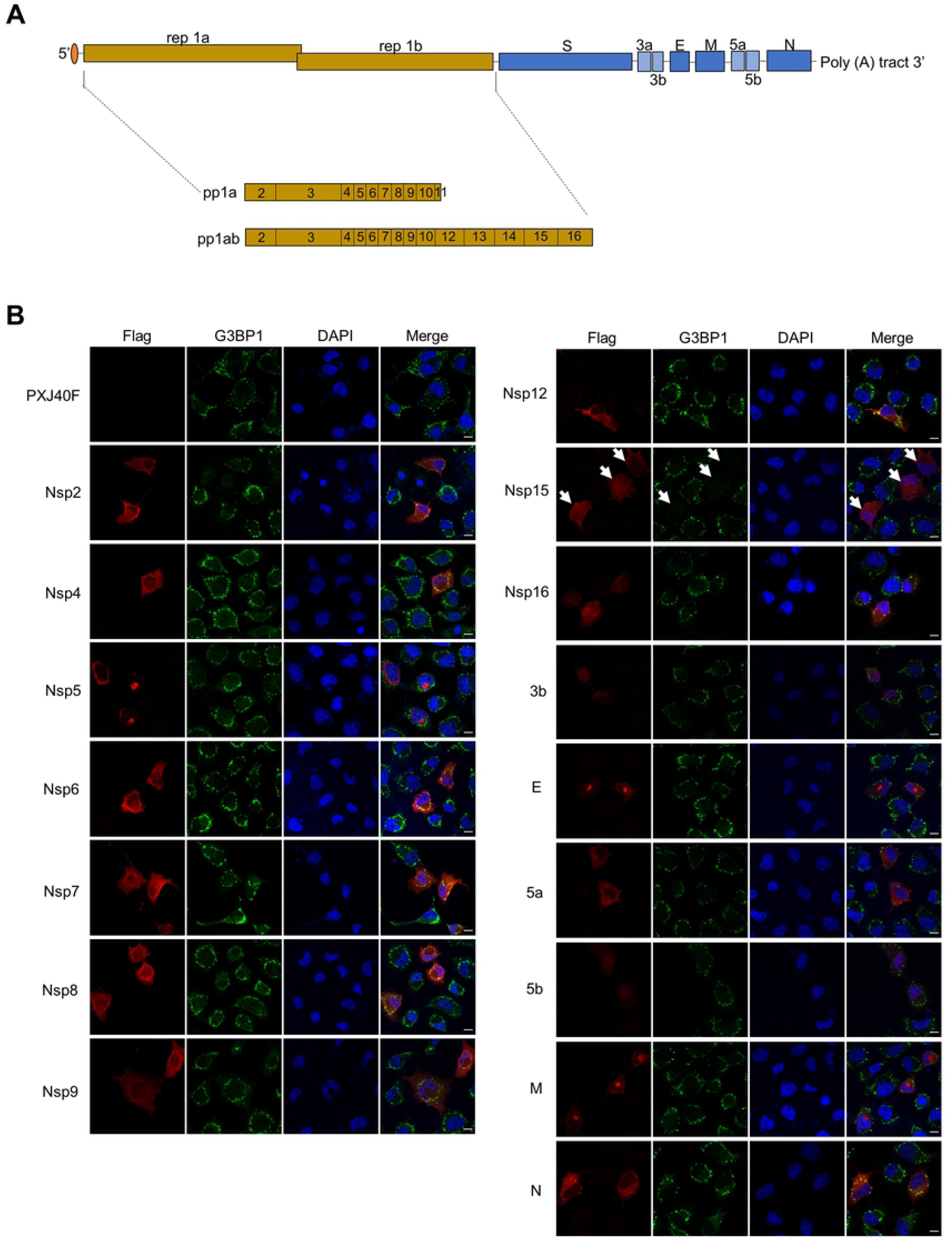
IBV nsp15 suppresses the formation of SGs. (A) Schematic diagram of the proteins encoded by IBV. (B) H1299 cells were transfected with plasmids encoding Flag-tagged IBV proteins or with vector PXJ40F. At 24 h post-transfection, cells received a heat shock treatment at 50°C for 20 min. IBV proteins were stained with anti-Flag antibody (red) and SGs were detected with anti-G3BP1 (green). Cell nuclei were stained with DAPI (blue). Shown are representative images out of three independent experiments. Scale bars: 10 μm.

Nsp15 is a conserved endoribonuclease of coronaviruses. It has been reported that its activity is involved in evasion of dsRNA sensing and interference with the type I IFN response [44, 50]. The conserved histidine (H) 223 and H238 of IBV nsp15 are critical for the endoribonuclease activity [51]. To examine whether the endoribonuclease activity is involved in the inhibition of SGs formation, we introduced an alanine (A) substitution in the catalytic core residues H223 or H238, to abrogate the catalytic activity. We next compared the ability of wild type nsp15 and mutant nsp15-H223A or nsp15-H238A to prevent SGs formation in H1299 cells. As expected, wild type nsp15 blocked the formation of SGs induced by heat shock, sodium arsenite, or NaCl (indicated with white arrow), while nsp15-H223A and nsp15-H238A did not (Fig 5A-C). Thus, the nsp15 endoribonuclease activity is required for the suppression of eIF2α-dependent and -independent SGs formation.

**Fig 5.**
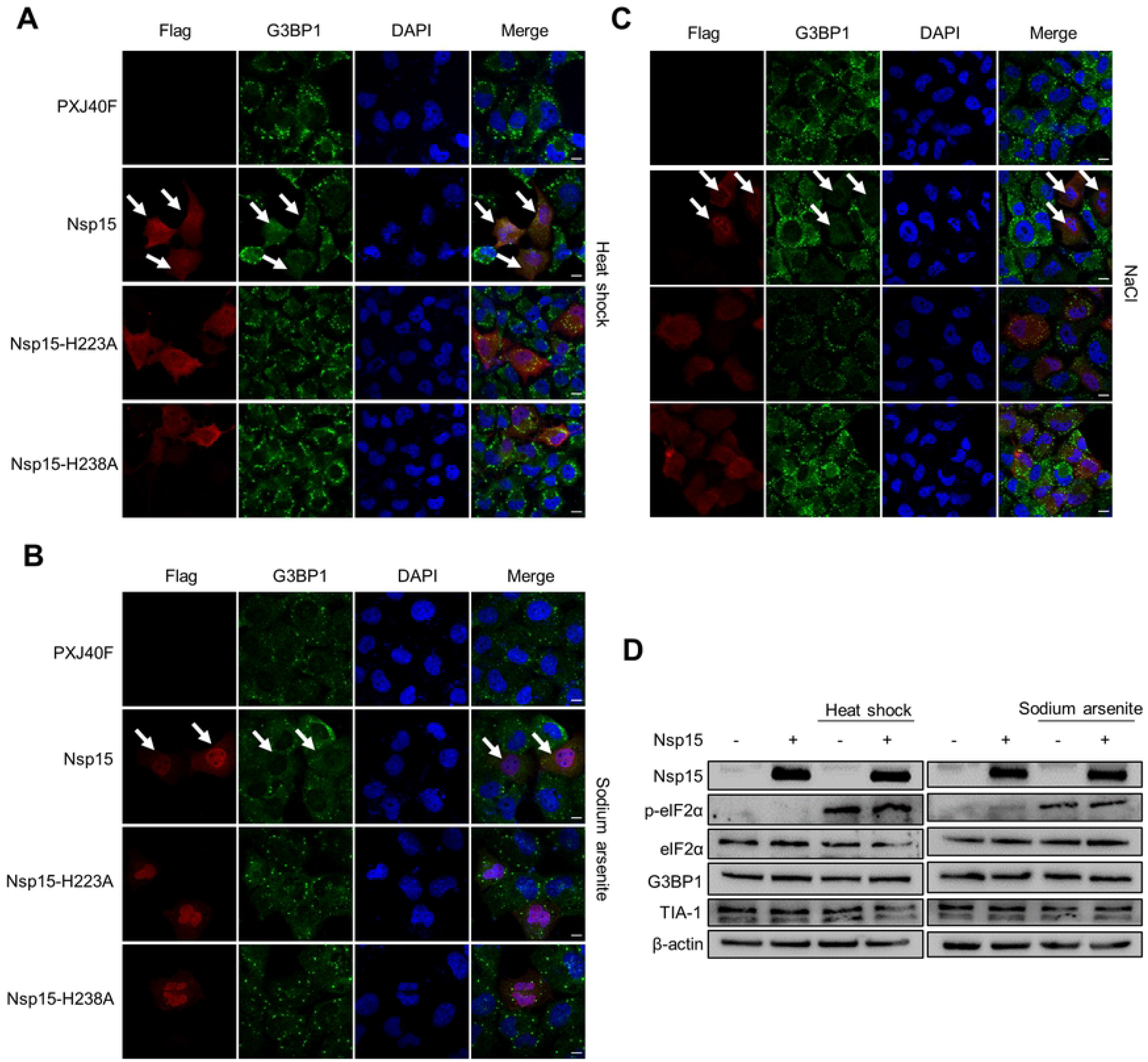
IBV nsp15 endoribonuclease activity is required for suppression of eIF2α-dependent and -independent formation of SGs. (A-C) H1299 cells were transfected with plasmids encoding IBV nsp15, nsp15-H223A, or nsp15-H238A, respectively. At 24 h post-transfection, cells received a heat shock treatment (50°C for 20 min), or 1 mM sodium arsenite for 30 min, or 200 mM NaCl for 50 min. Nsp15, nsp15-H223A, and nsp15-H238A were detected with anti-Flag antibody (red) and G3BP1 was detected with anti-G3BP1 (green). Cell nuclei were stained with DAPI (blue). Shown are representative images out of three independent experiments. Scale bars: 10 μm. (D) H1299 cells were transfected with plasmids encoding IBV nsp15 or with vector PXJ40F. At 24 h post-transfection, cells received a heat shock treatment (50°C for 20 min), or 1 mM sodium arsenite for 30 min. Cell lysates (10 μg/lane) were subjected to Western blot analysis, to check the expression of Flag-nsp15 and to determine the levels of phospho-eIF2α, eIF2α, G3BP1, TIA-1, and β-actin. Shown are representative bots out of three independent experiments.

In a preliminary investigation of how nsp15 may prevent SG formation, we examined whether nsp15 interferes with the phosphorylation of eIF2α. We observed a significant increase of phospho-eIF2α by sodium arsenite or heat shock treatment (Fig 3D-E); however, in agreement with the Fig 3D-E data on IBV-infected cells, no reduction of phospho-eIF2α was observed in nsp15-expressing cells (Fig 5D). No difference was observed also in the protein levels of G3BP1 and TIA-1 in nsp15-expressing cells, compared to control cells. Taken together, these data indicate that nsp15 interferes with the formation of SGs downstream of eIF2α.

### Nsp15-defective rIBV-nsp15-H238A induces canonical SGs by accumulation of dsRNA and activation of PKR

To further confirm the involvement of the endoribonuclease activity of nsp15 in the disruption of SGs formation, we constructed nsp15-defective recombinant virus rIBV-nsp15-H238A in which the nsp15 catalytic site H238 was replaced with an alanine (Fig 6A). We also constructed, but failed to recover nsp15-defective rIBV-nsp15-H223A, possibly due to the effect of the disruption of the catalytic site on virus replication. We compared the ability of wild type IBV and rIBV-nsp15-H238A to induce the formation of SGs in H1299 and DF-1 cells. At 20 h.p.i., only 18% of H1299 cells and 17% of DF-1 cells infected with wild type IBV showed the presence of SGs, whereas approximately 78% of the H1299 cells and 75% of DF-1 cells infected with rIBV-nsp15-H238A showed SGs formation (Fig 6B). Treatment with cycloheximide (CHX), a chemical which disassembles *bona fide* SGs, dissolved the rIBV-nsp15-H238A-induced G3BP1 and G3BP2 granules (Fig 6C), confirming that rIBV-nsp15-H238A induces canonical SGs.

**Fig 6.**
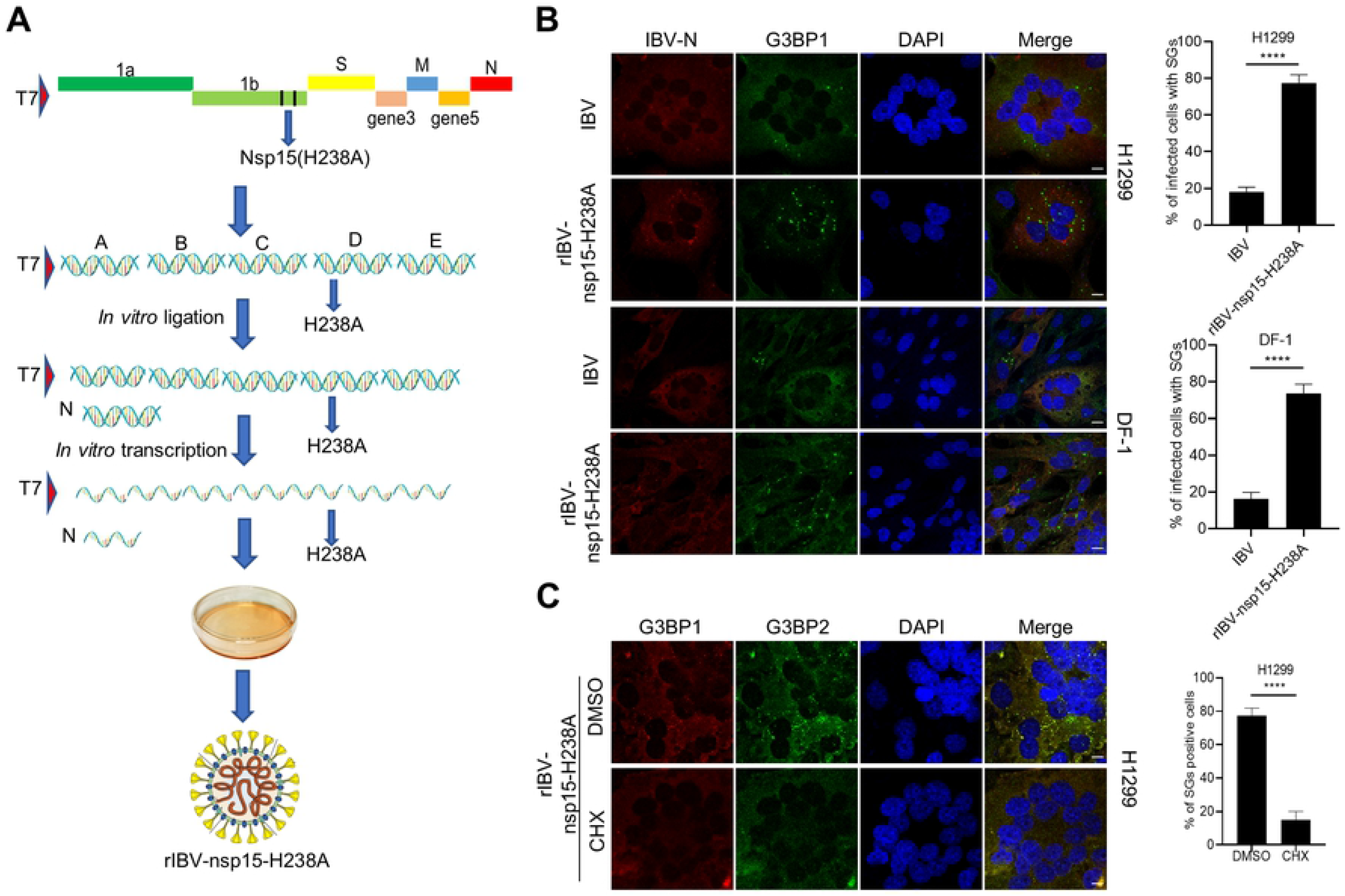
Nsp15-defective virus rIBV-nsp15-H238A induces canonical SGs by activation of PKR. (A) Schematic diagram of the construction of rIBV-nsp15-H238A as described in the methods section. (B) H1299 cells and DF-1cells were infected with IBV or rIBV-nsp15-H238A at a MOI of 1 for 20 h followed by immunostaining. Infected cells were identified with anti-IBN-N (red) and the SGs with anti-G3BP1 (green) antibody. Cell nuclei were stained with DAPI (blue). The bar graph on the right side indicate the mean + SD of the percentage of infected cells showing the presence of SGs was calculated over 20 random fields acquired for each condition. Values are representative of one out of three independent experiments. (C) H1299 cells were infected with rIBV-nsp15-H238A for 20 h and treated with 100 μg/ml of CHX for 1 h or with an equivalent volume of DMSO as control, followed by immunostaining with anti-G3BP1 or anti-G3BP2 antibodies. The bar graph on the right shows the percentage of SGs positive cells out of the total cells imaged. Values are representative of one out of three independent experiments. *P* values were calculated by Student’s test. ****, *P* < 0.0001 (highly significant). Scale bars: 10 μm.

In agreement, rIBV-nsp15-H238A significantly activated PKR by phosphorylation and in turn phosphorylated eIF2α, while wild type IBV did not (Fig 7A). Thus, nsp15 endoribonuclease activity is involved in antagonizing PKR activation, the well characterized dsRNA sensor and IFN-β inducer. We noted that the replication of rIBV-nsp15-H238A was impaired, as evidenced by the decreased level of IBV-S, IBV-M, and IBV-N protein synthesis, compared to wild type IBV (Fig 7A). Moreover, although rIBV-nsp15-H238A replication was low, it significantly stimulated the transcription of IFN-β at 20 h.p.i., which was approximately 25-fold higher than that induced by wild type IBV in H1299 cells (Fig 7B, left panel), and approximately 380-fold higher than that by wild type IBV in DF-1 cells (Fig 7B, right panel). Taken together, the activation of PKR by rIBV-nsp15-H238A and associated induction of type I IFN might be responsible for the lower replication of this recombinant virus.

**Fig 7.**
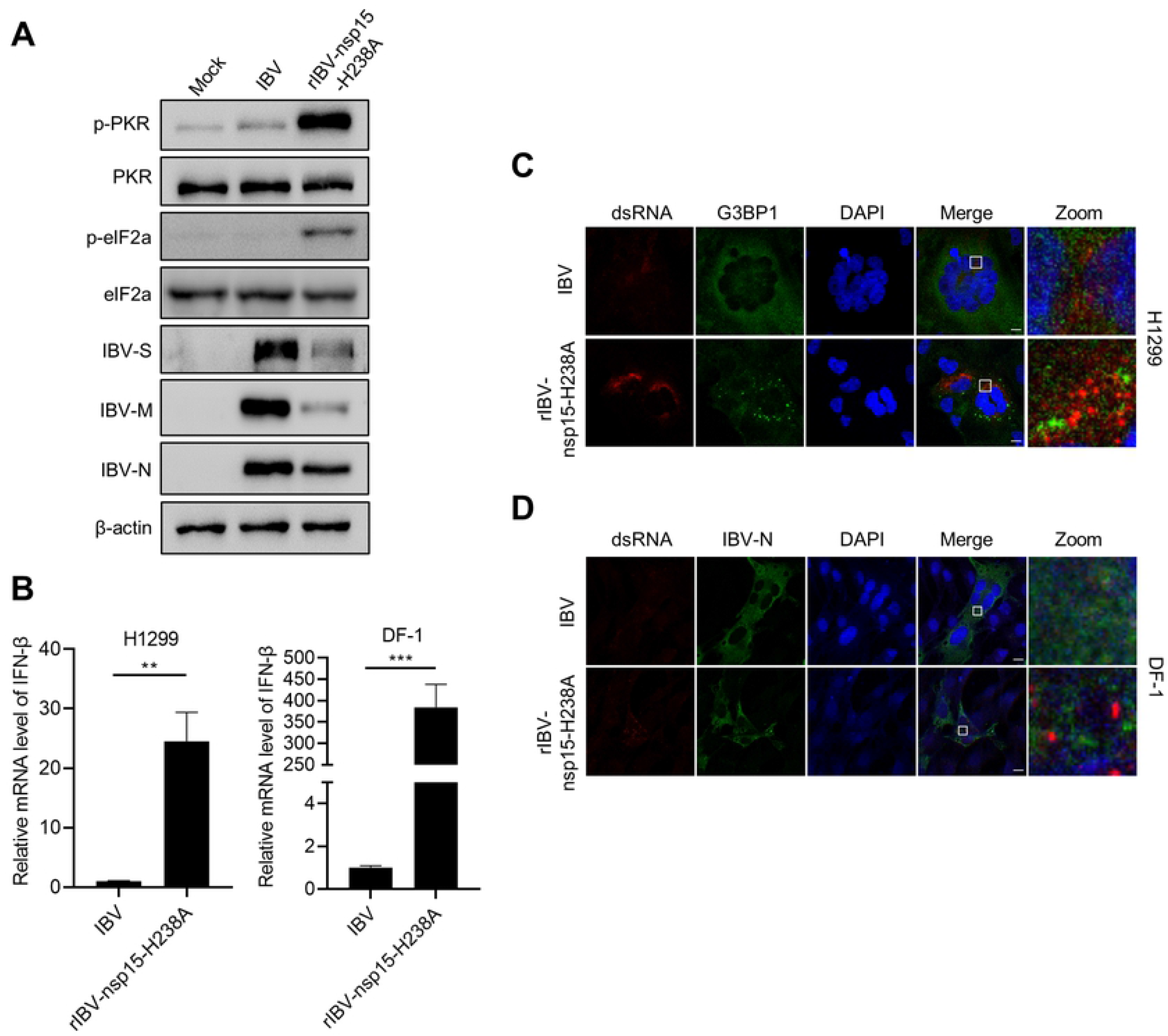
Nsp15-defective virus rIBV-nsp15-H238A strongly activates PKR by promoting dsRNA accumulation and eventually stimulates IFN response. (A) H1299 cells were mock infected or infected with IBV or rIBV-nsp15-H238A of 1 MOI for 20 h. Cells were lysed for western blotting analysis to detect the level of p-PKR, PKR, p-eIF2α, eIF2α, IBV-S, IBV-M, IBV-N, and β-actin. (B) H1299 and DF-1 cells were infected with IBV or rIBV-nsp15-H238A for 20 h respectively. Total RNA was extracted and subjected to quantitative RT-PCR to determine the transcription of *IFN-β*. Values are representative of three independent experiments. *P* values were calculated by Student’s test. **, *P* < 0.01; ***, *P* < 0.001. (C-D) H1299 and DF-1 cells were infected with IBV or rIBV-nsp15-H238A for 20 h followed by immunostaining. dsRNA (red) was detected with J2 antibody and G3BP1 or IBV-N (green) were determined with corresponding antibodies. Images shown are representative of three independent experiments. Scale bars: 10 μm.

The activation of PKR by rIBV-nsp15-H238A prompted us to measure and compare the levels of dsRNA during infection by using the specific J2 monoclonal antibody, which binds dsRNA greater than 40 nucleotides in length [51] and was previously successfully used during IBV infection in chicken cells [52]. Immunofluorescence analysis at 20 h.p.i., revealed evident accumulation of dsRNA in rIBV-nsp15-H238A infected H1299 cells, compared to wild type IBV infected cells (Fig 7C). The dsRNA produced by rIBV-nsp15-H238A however, did not colocalized with G3BP1 granules, suggesting the dsRNA is not recruited to SGs (Fig 7C). In DF-1 cells, we also observed higher levels of dsRNA accumulation during infection with rIBV-nsp15-H238A than with wild type IBV (Fig 7D). dsRNA dot blot analysis also supported the observation that infection with rIBV-nsp15-H238A leads to higher accumulation of dsRNA than with wild type IBV (data not shown). Although the dsRNA in wild type IBV infected cells partially co-localized with IBV-N, the dsRNA produced by rIBV-nsp15-H238A did not co-localize well with IBV-N (Fig 7C-D). We speculate that when compared to wild type IBV, rIBV-nsp15-H238A replication leads to higher accumulation of dsRNA and that the excess dsRNA may escape from replication-transcription complex (RTC). The “free” dsRNA in turn, triggers the activation of PKR and phosphorylation of eIF2α, results in translational shut off, eventually promoting the formation of SGs and activation of the type I IFN response. RT-PCR examination of the level of viral RNA showed that rIBV-nsp15-H238A indeed increases the ratio of negative strand RNA : positive strand RNA, compared to wild type IBV (supplementary Fig S2), suggesting the functional nsp15 is required for maintaining the ratio of viral (-:+) RNA. Altogether, these data indicate that intact nsp15 endoribonuclease activity acts to reduce the intracellular levels of dsRNA, thereby preventing activation of PKR and SGs formation.

### Nsp15-defective rIBV-nsp15-H238A strongly activates the IRF3-IFN signaling via the formation of SG

To examine the role of SGs in IBV infection, we knocked out the SGs core protein G3BP1 and G3BP2 in H1299 cells by a CRISPR-Cas9 approach. Depletion of G3BP1/2 resulted in the absence of SGs during sodium arsenite stimulation and rIBV-nsp15-H238A infection (Fig 8A). In G3BP1/2 knock out cells, the levels of phospho-TBK1 and phospho-IRF3 triggered by rIBV-nsp15-H238A infection were greatly decreased (Fig 8B); consequently, the transcription of *IFN-β* and ISG *IFIT1* induced by rIBV-nsp15-H238A infection were decreased (Fig 8C). These results suggest that the formation of SGs is necessary to elicit IRF3-IFN signaling in response to rIBV-nsp15-H238A infection. It was worth noting that rIBV-nsp15-H238A infection did not significantly stimulate p65 phosphorylation, and knock out of G3BP1/2 had no obvious effect on phospho-p65 levels (Fig 8B); thus, SGs formation is not involved in NF-κB signaling during IBV infection. Interestingly, upon G3BP1/2 knock out, we observed higher levels of IBV-S, IBV-M, and IBV-N (Fig 8B) and in agreement, more infectious progeny virus particles were produced, as evidenced by TCID_50_ assay (Fig 8D). This is in line with our previous suggestion that activation of the type I IFN signaling might be the factor limiting rIBV-nsp15-H238A replication; failure to trigger an IFN-β response in G3BP1/2 knock out cells however, promotes virus replication even in the absence of nsp15 endoribonuclease activity.

**Fig 8.**
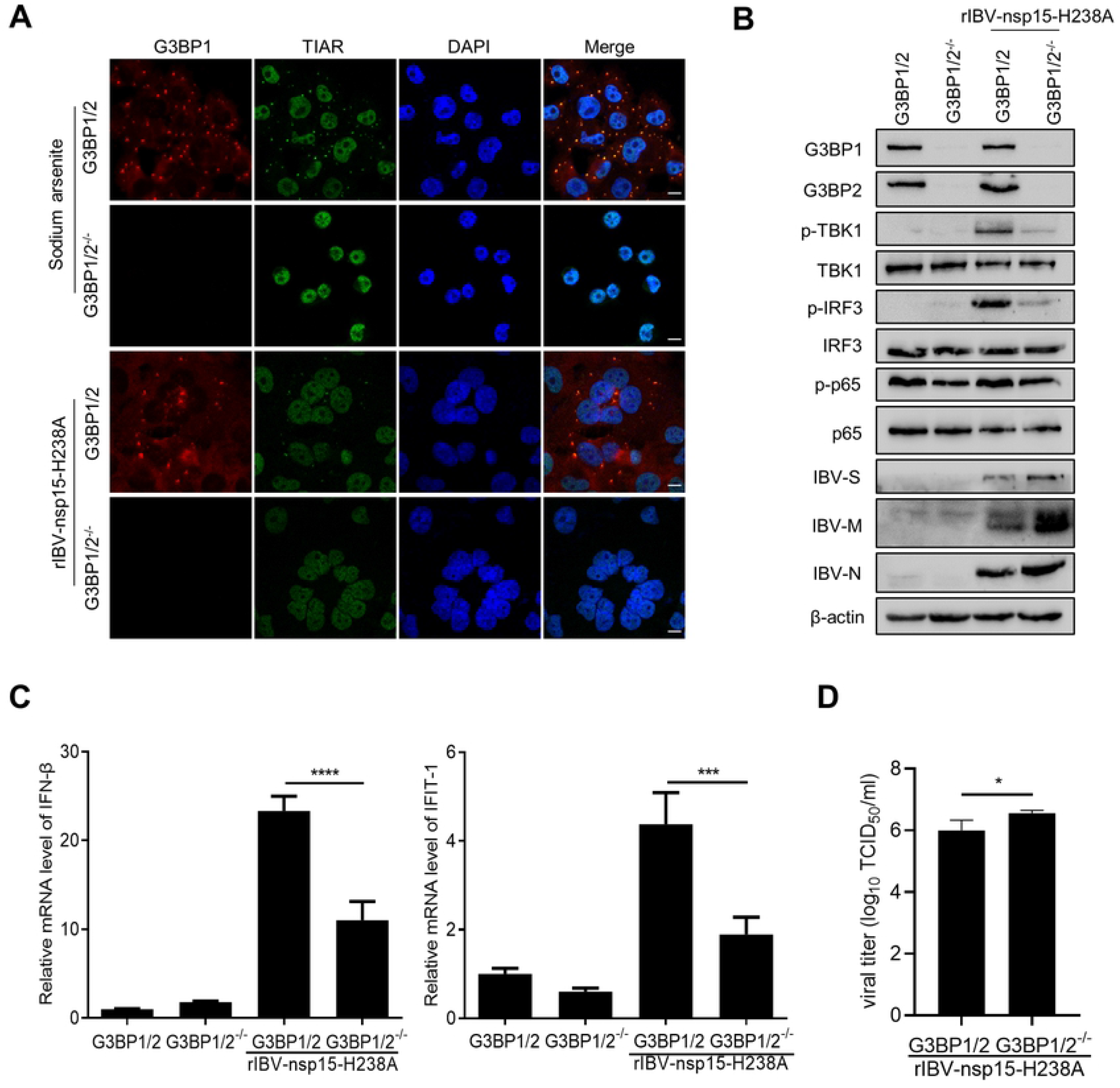
Depletion of SGs scaffold proteins reduces rIBV-nsp15-H238A induced IRF3-IFN-β signaling. (A) H1299 or H1299-G3BP1/2^-/-^ cells were treated with 1 mM sodium arsenite for 30 min or infected with rIBV-nsp15-H238A for 20 h, followed by immunostaining with anti-G3BP1 (red) and anti-TIAR (green). (B) H1299 cells and H1299-G3BP1/2^-/-^ cells were mock infected or infected with rIBV-nsp15-H238A for 20 h. Cell lysates were analyzed by western blot to detect G3BP1, G3BP2, p-TBK1, TBK1, p-IRF3, IRF3, p-p65, p65, IBV-S, IBV-M, IBV-N, and actin. (C) H1299 cells and H1299-G3BP1/2^-/-^ cells were inoculated with rIBV-nsp15-H238A for 20 h and the induction of *IFN-β* and *IFIT1* was quantified by quantitative RT-PCR. The bar graph shows means ± SD of the levels of *IFN-β* or *IFIT1*. (D) The supernatant from rIBV-nsp15-H238A infected H1299 and H1299-G3BP1/2^-/-^ cells was collected at 20 h.p.i. and virus titers (TCID_50_) calculated. The bar graph shows means ± SD of three independent determination of viral titer. Data are representative of three independent experiments. *, *P* < 0.1; **, *P* < 0.01.

To investigate whether the involvement of SGs in IRF3-IFN signaling is restricted to specific virus infections, we examined the IRF3-IFN signaling upon poly I:C stimulation. Results showed that in G3BP1/2 positive cells, poly I:C strongly stimulated phosphorylation of IRF3 and to a lesser extent of TBK1 (Fig 9A), and promoted IRF3 nuclear translocation (Fig 9B, 34% of total cells display nuclear IRF3); however, in the absence of G3BP1/2, poly I:C stimulation led to reduced TBK1 phosphorylation, and to a greater extent, to reduced IRF3 phosphorylation (Fig 9A), as well as less IRF3 nuclear translation (Fig 9B, 9% of total cells with nuclear IRF3). As a consequence, transcription of *IFN-β* and *IFIT-1* was significantly decreased upon poly I:C stimulation of G3BP1/2 knock out cells (Fig 9C). Altogether, these results demonstrate that SGs positively regulates IRF3-IFN signaling and that such a mechanism is not restricted to a specific virus infection.

**Fig 9.**
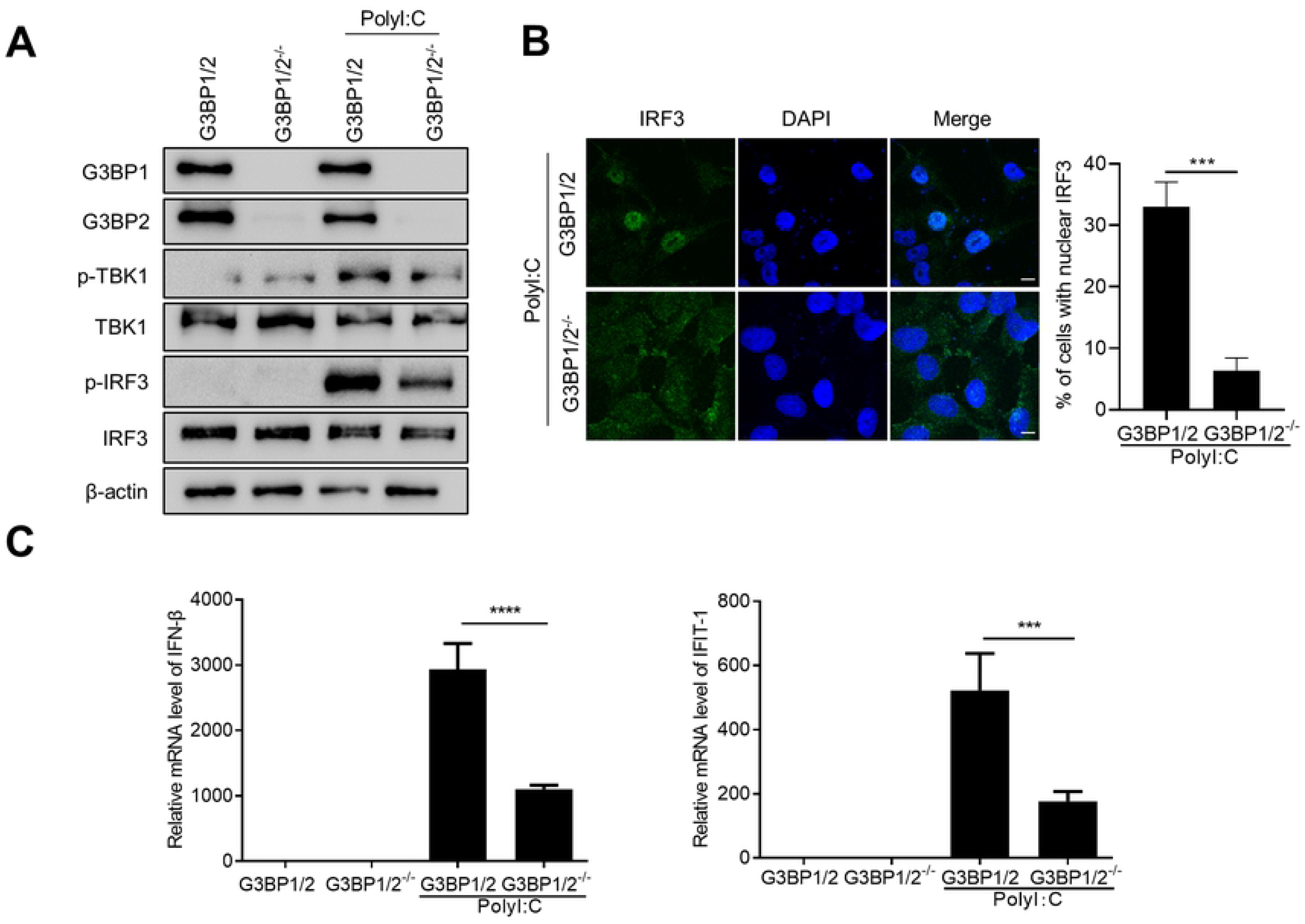
Depletion of SGs scaffold proteins reduces poly I:C induced IRF3-IFN-β signaling. (A-C) H1299 cells and H1299-G3BP1/2^-/-^ cells were transfected with poly I:C (1 μg/ml) for 6 h. The levels of G3BP1, G3BP2, p-TBK1, TBK1, p-IRF3, IRF3, and actin were determined by Western blot analysis (A), the nuclear translocation of IRF3 was examined by immunostaining (B), and the induction of *IFN-β* and *IFIT1* was quantified by quantitative RT-PCR (C). The bar graph shows means ± SD of the percentage of nuclear IRF3 positive cells out of the total cells imaged in (B) and the relative expression levels of *IFN-β* or *IFIT1* (C). Data are representative of three independent experiments. ***, *P* < 0.001; **, *P* < 0.01.

### Aggregation of PRRs and signaling intermediates to SGs during IBV infection

A previous report showed that the dsRNA sensors PKR, MDA5, and RIG-I are located to SG and sense dsRNA [24]. In this study, we examined the subcellular localization of PRRs and signaling intermediates during IBV infection. In the small proportion of IBV-infected cells that displayed the presence of SGs, PKR, MDA5, TLR3 and MAVS aggregated and colocalized with G3BP1 granules (Fig 10A). These results demonstrate that SGs indeed recruit PRRs and their signaling intermediates during IBV infection. We further examined the subcellular location of signaling intermediates, results showed that TRAF3, TRAF6, TBK1, and IKKε all aggregated to G3BP1 granules (Fig 10B). These data, combined with the positive role on IRF3-IFN signaling, demonstrate that SGs may function as a platform for PRRs and downstream signaling intermediates.

**Fig 10.**
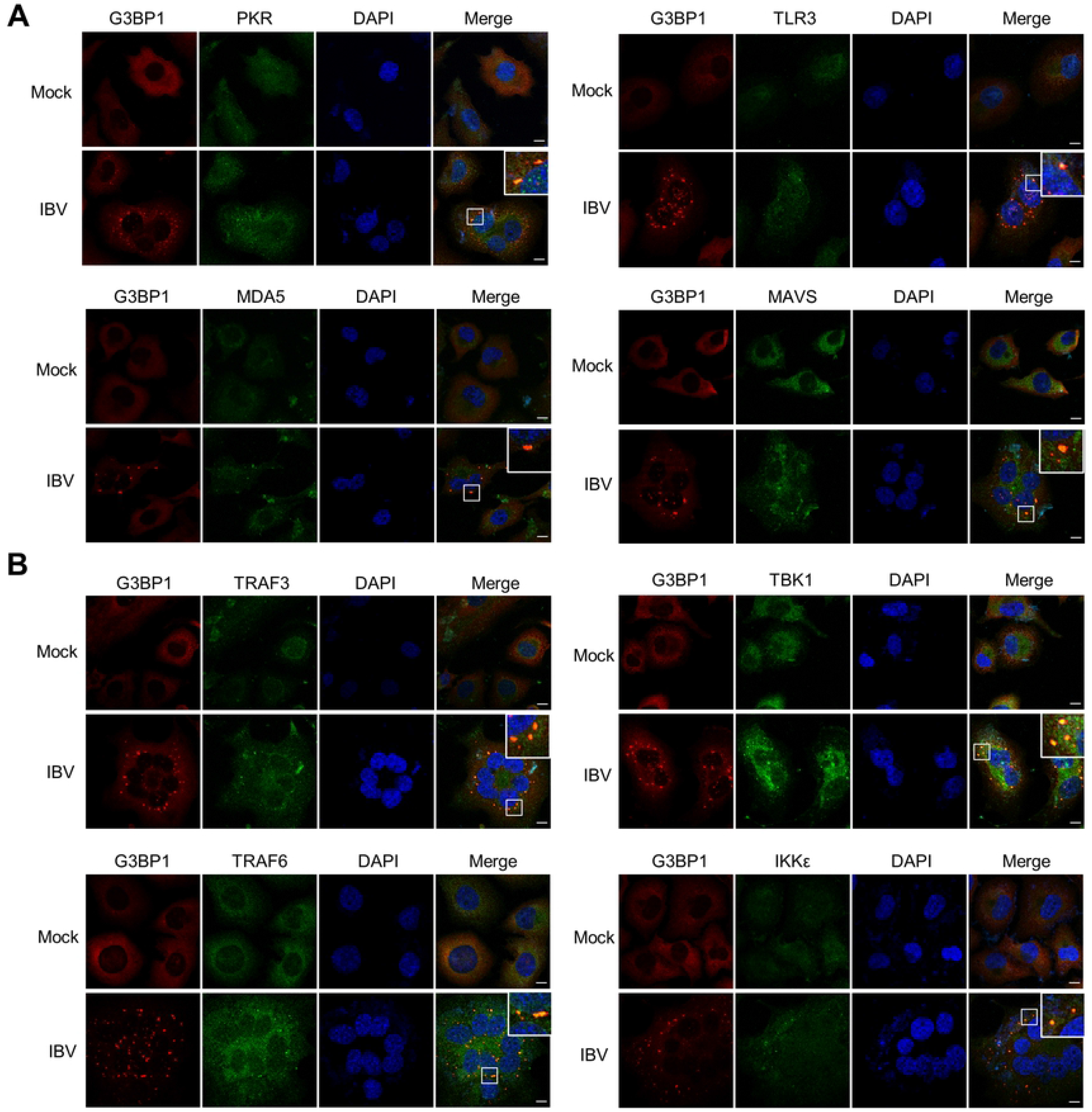
PRRs and innate immunity signaling intermediates aggregate to IBV-induced SGs. (A-B) H1299 cells were mock infected or infected with IBV followed by immunostaining at 20 h.p.i.. Anti-G3BP1 (red) was used to monitor SGs formation, and PKR, MDA5, TLR3, MAVS, TRAF3, TRAF6, TBK1, IKKε (green) were detected with corresponding antibodies. Cell nuclei were stained with DAPI (blue). Shown are representative images out of three independent experiments. Scale bars, 10 μm.

## Discussion

SGs formation or inhibition has been reported for different groups of coronaviruses: MHV and TGEV induce SGs or SG-like granules [34, 35], whereas MERS-CoV does not [32, 33], and IBV was reported to induce SGs formation but only in 20% of infected Vero cells [37], yet the biological significance of SGs in coronavirus replication is unclear. In this study, we report that IBV indeed induces SGs formation in a small proportion of infected cells and it does so not only in mammalian (H1299 and Vero) cells, but also in chicken DF-1 cells. Furthermore, consistent with previous reports [53], also in our study we found that IBV inhibits both eIF2α-dependent (heat shock, sodium arsenite) and -independent (NaCl) SGs formation. We also assessed SGs formation by porcine epidemic diarrhea virus (PEDV) infection, only 10%-20% of PEDV-infected Vero cells were SGs positive (data not shown). Combined with the inhibition of SGs formation by MERS-CoV [33], our results suggest that the inhibition of SG formation by coronavirus might be a universal phenomenon, not only restricted to a specific coronavirus. Screening of viral proteins involved in inhibition of SGs formation, revealed that nsp15 is a specific stress response antagonist: overexpression of nsp15 resulted in disruption of both eIF2α-dependent and -independent SGs formation, which could be attributed to its endoribonuclease activity; abrogating nsp15 endoribonuclease function *in vivo* led to impaired virus replication, efficient formation of SGs, accumulation of dsRNA, robust activation of PKR, and activation of IRF3-IFN signaling. Thus, functional nsp15 is specifically required for efficient virus replication as it plays a role in inhibition of SG formation and subsequent activation of an anti-viral response.

Coronaviruses are positive-stranded RNA viruses that replicate in the host cell cytoplasm. The viral RNA synthesis is performed in RTCs that include viral and cell proteins, connected with convoluted membranes and double membrane vesicles [54, 55]. Replication of the coronavirus genome requires continuous RNA synthesis, whereas transcription is a discontinuous process unique among RNA viruses. Transcription includes a template switch during the synthesis of sub-genomic negative strand RNAs to add a copy of the leader sequence [56–58]. The negative strand RNAs is the replication intermediate of genomic RNA and of sub-genomic RNA. During the replication and transcription process, positive and negative strand RNA form dsRNA. It has been reported that nsp13 and cellular helicases help to unwind the dsRNA for efficient replication and transcription. The amount of negative strand intermediates is approximately 10% of the positive strand RNA [59]. It is believed that the proper ratio of positive and negative stand RNA is important for efficient replication and transcription, as well as subsequent genome packaging and mRNA translation. How do coronaviruses modulate the ratio of positive and negative strand RNA? Coronavirus nsp15 has uridylate-specific endoribonuclease activity on single-stranded RNA and dsRNA [60, 61], is considered an integral component of the RTC and co-localizes with viral RNA [62]. It has been reported that nsp15 is involved in efficient viral RNA synthesis [63, 64]. Nsp15-null MHV exhibits severe replication defects in macrophages and is highly attenuated in mice [44, 50]. As the negative strand RNA intermediates harbor a poly (U) sequence, which is complementary to the positive strand RNA poly (A) tail, we propose that nsp15 targets negative strand intermediates or dsRNA intermediates within stalled RTCs that are no longer active in viral RNA synthesis. This is supported by the observation that, compared to wild type IBV, infection with nsp15-null virus rIBV-nsp15-H238A leads to substantial accumulation of dsRNA intermediates that do not localize with RTCs and to an increased ratio of negative strand:positive strand RNA. In this way, nsp15 controls viral RNA quantity for efficient replication/transcription, thereby facilitating the proliferation of virus. Recently, Hackbart reports that nsp15 cleaves the 5’-poly(U) from negative-sense viral RNA intermediates [65], confirming our hypothesis.

Previous reports show that coronavirus nsp15 and arterivirus nsp11 act as IFN antagonists [51]. Overexpression of SARS-CoV nsp15 inhibits the IFN response and MAVS-mediated apoptosis [66]. Interestingly, Hackbart find that the poly(U) containing negative-sense viral RNA is sufficient to stimulate MDA5, and nsp15 is responsible for the cleavage of the poly(U) containing negative-sense viral RNA, thereby antagonize the IFN response [65]. Here, we describe a previously unrecognized role of coronavirus nsp15 in the evasion of PKR activation and interference with SGs formation. RNA viruses that replicate via dsRNA intermediates can be detected as “non-self” by host dsRNA sensors: PKR, RIG-I, MDA5 and TLR3, eventually stimulating the production of type I IFN [19, 20, 22, 67]. It is likely that dsRNAs are shielded within double-membrane vesicles and replication intermediates are likely protected by the RTC and N protein. In this study, by generating nsp15-defective recombinant virus, we find that compared to the wild type virus, infection with rIBV-nsp15-H238A leads to dsRNA accumulation, PKR activation, robust formation of SG as well as up-regulation of *IFN-β*, which ultimately coincided with impaired rIBV-nsp15-H238A replication. Therefore, IBV nsp15 acts as IFN antagonist, likely through removal of dsRNA intermediates at sites of RNA synthesis, thereby efficiently evading integrated stress and innate anti-viral host responses. The involvement of the viral ribonuclease in degrading viral dsRNA and antagonizing IFN responses has also been reported in pestivirus and Lassa virus [68–70].

Several reports show that SGs serve as platform for viral dsRNA sensing by RLRs and subsequent activation of viral immune responses [71, 72]. Recent studies reported that several other IFN regulatory molecules, such as MEX3C, Riplet, DHX36 and Pumilios, also localize to SGs [73]. It is thus reasonable that viruses evolved mechanisms to suppress SGs formation in order to promote their propagation. Influenza A virus (IAV) non-structural protein 1 (NS1) is reported to be involved in subversion of PKR-dependent SG formation [25]; importantly, during NS1-null IAV infection, viral RNAs and nucleocapsid protein co-localize in SGs together with RIG-I, PKR and SGs markers G3BP1/TIAR; knock down of the G3BP1 or PKR genes abrogated NS1-null IAV-induced IFN production, concomitantly with defects in SGs formation. Here, we find that wild type IBV triggers the formation of SGs only in 20% infected cells, and that PRRs (PKR, MDA5, TLR3) and signaling intermediates (MAVS, TRAF3, TRAF6, TBK1, IKKε) aggregate to the IBV-induced SGs. Nsp15-null recombinant IBV robustly activates PKR, efficiently induces SGs formation (80% infected cells with SGs formation), and strongly induced the transcription of *IFN-β;* but in SGs’ core proteins defective cells, either by nsp15-null recombinant IBV infection or poly I:C stimulation, the induction of *IFN-β* signaling is severely impaired. These data thus further confirm that SGs play a positive regulatory role in the IRF3-IFN signaling, leading to the initiation of anti-viral innate responses, and that this is not restricted to a specific virus infection. These observations strongly suggest that the formation of SGs is critical for virus-induced antiviral innate immunity and SGs may function as a scaffold for viral RNA recognition by RLRs.

Virus-encoded endoribonucleases not only modulate viral RNA, but also target the majority of cellular mRNAs, likely enabling viral mRNAs to better compete for limiting translation components and directing the cell from host to virus gene expression. Targeting host mRNA for degradation not only restricts host gene expression, but also subverts SGs by depleting the core component of SG, RNA. Thus, the ribonuclease is a unique strategy for viruses to subvert SGs. It has been well characterized that the herpes simplex virus 1 (HSV-1) and HSV2 employ the virion host shutoff (VHS) endoribonuclease to impede the SGs formation [74, 75]. Infection with a mutant virus lacking VHS (ΔVHS) results in PKR activation and PKR-dependent SGs formation in multiple cell types [76, 77]. Destabilization of host mRNAs by VHS may directly contribute to its disruption of SG formation [78]. In addition to be important in virus replication, coronavirus nsp15 may also target host mRNA and subsequently inhibit host protein translation. Interestingly, we previously showed that differently from alphacoronaviruses such as MHV and SARS-CoV [79, 80], wild-type IBV affects host protein synthesis leading to host-protein shut off, but without affecting mRNA stability [81]. In the current study, when nsp15 was overexpressed, it was worth noting that nsp15 located to nucleus (Fig 4, Fig 5A-C). Taken all our observations together, we speculate that nuclear nsp15 may interfere with the host pre-mRNA processing or nuclear export. Further studies are needed to fully elucidate the mechanisms used by nsp15 to target to host protein expression system.

Altogether, there are several possibilities that may account for the lack of SGs formation upon transfection of nsp15 or infection with IBV (Fig 11): (1) as SGs are dynamic foci, nsp15 might be involved in the disassembly of SGs. The requirement for nsp15 endoribonuclease activity for the disruption of both eIF2α-dependent or eIF2α-independent SGs, implies that removal of mRNA from SGs promotes their disassembly and predicts that intact RNA is crucial for maintaining the integrity of SGs; (2) during IBV infection, nsp15 endoribonuclease activity may be involved in the regulation of virus genomic RNA replication and sub-genomic mRNA transcription, consequently functioning as a “gatekeeper” to sequester viral dsRNA within replication complexes and away from host sensor PKR, resulting in absence of SG formation; (3) nsp15 prevents the assembly of SGs by promoting the destruction of mRNAs present in polysome, free mRNA, or by blocking the processing of pre-mRNA and nuclear export of mRNA, thus preventing a crucial step in SGs assembly pathway. Our unpublished data show that nsp15 interferes with the host protein translation and retains poly(A) binding protein 1 (PABP1) in the nucleus, and that the endoribonuclease activity is specifically required for this function. Thus, nsp15 probably also targets host mRNA to prevent SGs assembly. As the nsp15 endoribonuclease is conserved, we speculate that coronaviruses employ similar mechanisms to antagonize the host anti-viral SGs formation for efficient virus proliferation. Altogether, this study is the first to demonstrated that IBV coronavirus antagonize the formation of antiviral hubs SGs through the activity of its endoribonuclease nsp15, and that this is required for efficient virus proliferation.

**Fig 11.**
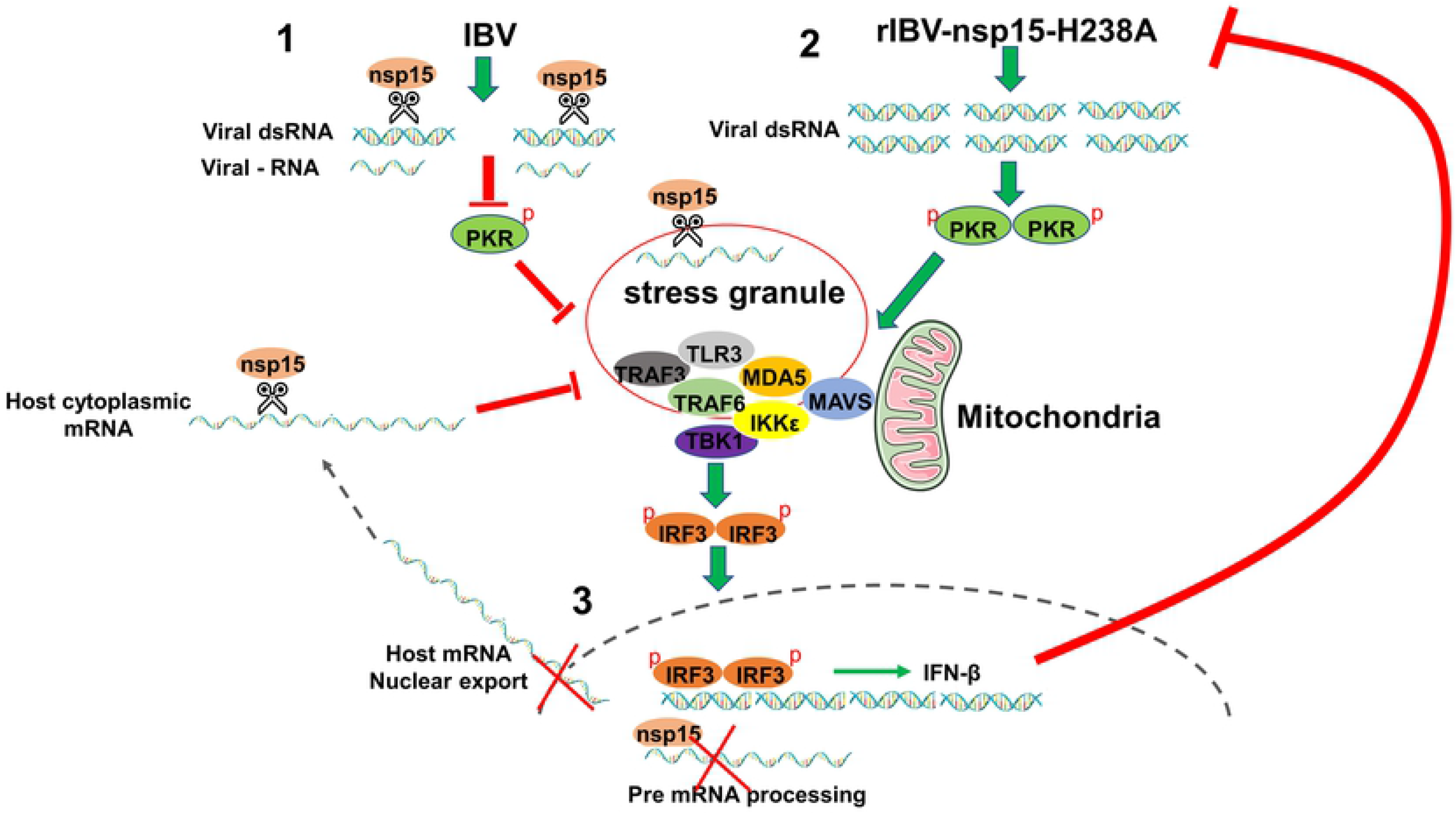
The working model of inhibition of anti-viral stress granule formation by infectious bronchitis virus nsp15. 1) IBV genome replication and mRNA transcription produce dsRNA. Nsp15 functions to cleave viral dsRNA and reduce its accumulation. Thus, virus avoids activation of PKR and impedes SGs formation. 2) Absence of nsp15 endoribonuclease activity in rIBV-nsp15-H238A, results in the accumulation of viral dsRNA, activation of PKR, and subsequent formation of SGs. The aggregation of PRRs and signaling intermediates to SGs facilitates the signaling transduction and IRF3 activation, finally inducing the expression of *IFN-β*. Production of IFN-β in turn, effectively limits rIBV-nsp15-H238A proliferation. 3) In parallel, nsp15 may also cleave host mRNA or interfere with host mRNA nuclear export or processing thereby preventing the assembly of SGs or promoting their disassembly.

## Materials and methods

### Cells and viruses

H1299 cells were purchased from Cell Bank of China Academy of Science and were maintained in Roswell Park Memorial Institute (RPMI) 1640 medium supplemented with 10% (v/v) fetal calf serum (FCS). Vero and DF-1 cells were purchased from ATCC and were grown in Dulbeco’s modified eagle medium (DMEM) with 10% FCS. IBV Beaudette strain was a gift from Prof Dingxiang Liu’s lab, South China Agricultural University. The recombinant virus IBV-nsp15-H238A was constructed in our laboratory with the technical support of Prof. Shouguo Fang, Yangtze University, as further detailed below.

### Antibodies and chemicals

Rabbit anti-IBV-S, rabbit anti-IBV-M, rabbit anti-IBV-N and rabbit anti-nsp3 were the gifts from Prof Dingxiang Liu’s lab, South China Agricultural University. Below we provide the list of all primary antibodies used, all of them directed against mammalian proteins; their dilution and eventual cross-reactivity to chicken proteins of interest is summarized in Table 1. Rabbit anti-G3BP1 (ab181150), rabbit anti-G3BP2 (ab86135), rabbit anti-phospho-PKR (ab32036), rabbit anti-phospho-IRF3 (ab76493), and mouse anti-G3BP1 (ab56574) were purchased from Abcam; rabbit anti-TIAR (#8509), rabbit anti-PKR (#12297), rabbit anti-eIF2α (#5324), rabbit anti-phospho-eIF2α (#3398), rabbit anti-MDA5 (#5321), rabbit anti-TLR3 (#6961), rabbit anti-MAVS (#24930), rabbit anti-TARF3 (#61095), rabbit anti-TRAF6 (#8028), rabbit anti-IKKε (#3416), rabbit anti-TBK1 (#3504), rabbit anti-phospho-TBK1 (#5483), rabbit anti-IRF3 (#11904), rabbit anti-p65 (#8242), and rabbit anti-phospho-p65 (#3033) were purchased from Cell Signaling Technology; mouse anti-TIA-1 (sc-116247) was purchased from Santa Cruz; mouse anti-Flag (F1804) was purchased from Sigma; rabbit anti-β-actin (AC026), goat anti-rabbit IgG (H+L) (AS014), and goat anti-mouse IgG (H+L) (AS003) conjugated with HRP were from Abclonal; J2 mouse anti-dsRNA (10010200) was purchased from Scicons. Alexa Fluor goat anti-rabbit-488 (A-11034), Alexa Fluor goat anti-rabbit-594 (A-11037), Alexa Fluor goat anti-mouse-488 (A-11029), and Alexa Fluor goat anti-mouse-594 (A-11005) were obtained from Invitrogen. Sodium arsenite (S7400) was purchased from Merck. Poly I:C (31852-29-6) was from InvivoGen.

### Plasmids construction and transfection

The plasmids encoding IBV nsp2, nsp4, nsp5, nsp6, nsp7, nsp8, nsp9, nsp12, nsp15, nsp16, 3b, E, 5a, 5b, M and N were generated by amplification of cDNA from IBV Beaudette-infected Vero cells using corresponding primers (Table 2) and cloned into PXJ40F. The restriction endonuclease sites for most inserts were *BamH I* and *Xho I*, while the restriction endonuclease sites for M are *EcoR I* and *Xho I*. The catalytic mutant plasmids of IBV nsp15 was cloned by using Mut Express II Fast Mutagenesis Kit V2 (Vazyme) as further detailed below. The mutagenesis primers are shown in table 2.

**Table 2.**
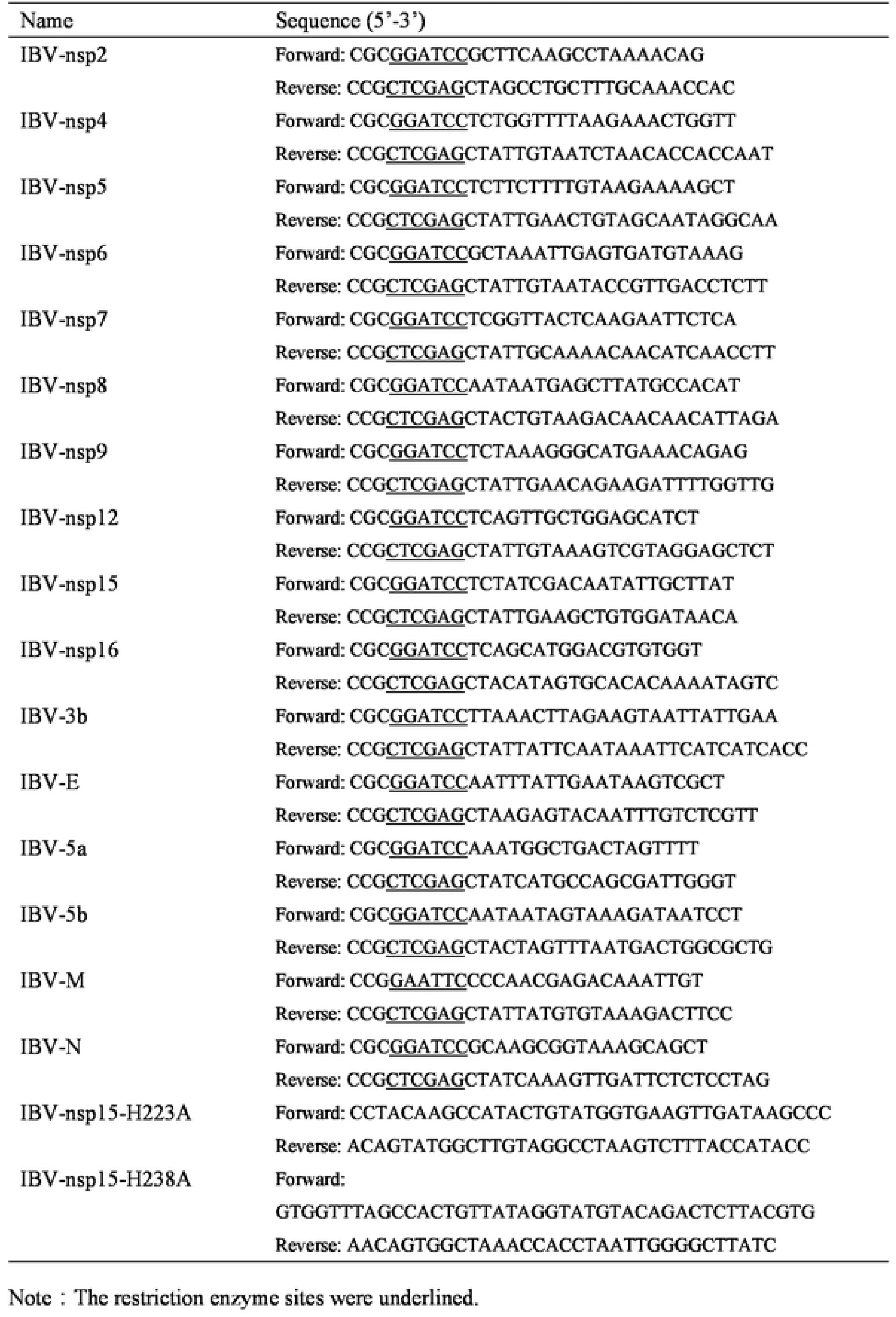
Primers used to construct plasmids

Cells were seeded on glass coverslips in a 24 wells cluster (25,000 cells/well). The indicated plasmids were transfected into cells using Fugene HD (Promega) according to the manufacturer’s handbook. Briefly, 0.5 μg plasmid and 1.5 μl Fugene HD (m/v=1:3) were diluted and incubated in 0.25 ml OptiMEM (Gibco). After 5 min, plasmid and Fugene HD were mixed and incubated at room temperature for 15 min, allowing the formation of lipid-plasmid complex. Finally, the complex was added to the cultured cells and incubated for 24 h.

### Indirect immunofluorescence and confocal microscopy

Cells were seeded on glass coverslips in a 24 wells cluster (25,000 cells/well) and the next day were infected with virus or transfected with various plasmids or with poly I:C (0.25 μg/well). At the indicated time points, cells were treated with heat shock (50°C, 20 min), sodium arsenite (1 mM, 30 min), NaCl (200 nM, 50 min), or cycloheximide (CHX-100 μg/ml, 1 h), in the latter case DMSO was used as negative control. Cells were then fixed with 4% paraformaldehyde in PBS for 15 min at room temperature. After three washes with PBS, cells were permeabilized with 0.5% Triton X-100 in PBS for 15 min and incubated in blocking buffer (3% BSA in PBS) for 1 h. Cells were incubated with the primary antibody diluted in blocking buffer (as indicated in Table 1) overnight at 4°C, followed by incubation with Alexa Fluor conjugated secondary antibody diluted with 1:500 in blocking buffer for 1 h at 37°C. In case of double staining, cells were incubated with a different unconjugated primary antibody, followed by incubation with the corresponding conjugated secondary antibody and incubated as described before. Between and after each incubation step, the cell monolayer was washed three times with blocking buffer. DAPI was then applied to stain nuclei for 15 min. Finally, cells were washed once with PBS and examined by Zeiss LSM880 confocal microscope.

### Quantitative RT-PCR analysis

Total cellular RNAs were extracted using Trizol reagent (Ambion). cDNAs were synthesized from 2μg total RNA using oligo(dT) primers and M-MLV reverse transcriptase system (Promega). cDNA was used as template for quantitative PCR using a Bio-Rad CFX-96 real time PCR apparatus and SYBR green master mix (Dongsheng Biotech). PCR conditions were as follow: an initial denaturation at 94°C for 3 min, 40 cycles of 94°C for 15 s, 60°C for 15 s and 72°C for 20 s. The specificity of the amplified PCR products was confirmed by melting curve analysis after each reaction. The primers used were: for human *IFN-β*, 5’-GCTTGGATTCCTACAAAGAAGCA-3’ (F) and 5’-ATAGATGGTCAATGCGGCGTC-3’ (R); for human *IFIT1*, 5’-GCCATTTTCTT TGCTTCCCCT-3’ (F) and 5’-TGCCCTTTTGTAGCCTCC TTG-3’ (R); for human *β-actin*, 5’-GATCTGGCACCACACCTTCT-3’ (F) and 5’-GGGGTGTTGAAGGTC TCAAA-3’ (R); for chicken *β-actin*, 5’-CCAGACATCAGGGTGTGATGG-3’ (F) and 5’-CTCCATATCATCCCAGTTGGTGA-3’ (R); for chicken *IFN-β*, 5’-GCTCTCACCACCACCTTCTC-3’ (F) and 5’-GCTTGCTTCTTGTCCTTGCT-3’ (R); for IBV positive strand RNA: 5’-GTCTATCGCCAGGGAAATGTCT-3’ (F) and 5’-GTCCTAGTGCTGTACCCTCG-3’(R), which target to 3’ untranslated region of virus genome; for IBV negative strand RNA: 5’-GTCCTAGTGCTGTACCCTCG-3’ (F) and 5’-GTCTATCGCCAGGGAAATGTCT-3’(R), which target to 5’ sequence of virus negative strand RNA. The relative expression of each gene or virus RNA was normalized to *β-actin* mRNA levels and calculated using the 2^−ΔΔCT^ method. All assays were performed in triplicate and the results are expressed as the means ± standard deviations.

### Western blotting analysis

Cells were lysed in 2x protein loading buffer (20 mM Tris-HCl, 2% SDS, 100 mM DTT, 20% glycerol, 0.016% bromophenol blue). Cell debris was pelleted at 15000 × g for 10 min and 10 μg of the cleared cell lysates were resolved on a 10% SDS-PAGE and transferred to 0.45 μm nitrocellulose membrane (GE life Sciences). Membranes were blocked in blocking buffer (5% non-fat milk, TBS, 0.1% Tween 20) for 1 h, followed by incubation with primary antibody diluted in blocking buffer as indicated in S2 table overnight at 4°C. The membranes were then incubated with secondary antibodies diluted in blocking buffer as indicated in S2 table for 1 h at room temperature. Between and after the incubations, membranes were washed three time with washing buffer (0.1% Tween in TBS). The signals were developed with luminol chemiluminescence reagent kit (Share-bio) and detected using Tanon 4600 Chemiluminescent Imaging System (Bio Tanon).

### Quantification of stress granules formation and IRF3 nuclear translocation in viral infected cells

For quantification of SGs formation, images from 20 random high-powered fields were captured. The number of infected cells (IBV-N positive) in the acquired fields was counted. Cells displaying IBV-N expression and G3BP1 foci were counted as positive for SGs formation. The relative percentage of infected cells showing SGs formation was calculated as: (number of cells with G3BP1 granules and IBV-N protein expression divided by the total number of IBV-N positive cells) x 100. Similarly, for quantification of nuclear IRF3, 20 random high-powered fields were captured, and the percentage of cells displaying nuclear IRF3 out of all imaged cells was calculated.

### Generation of G3BP1/2 knock out cell

Lenti CRISPRv2 was ligated with a pair of guide sequences targeting G3BP1/2 exon 1 which were designed by Zhang’lab (https://zlab.bio/guide-design-resources). The sgRNA of G3BP1 is 5’-CACCGTGTCCGTAGACTGCATCTGC-3’ and G3BP2 is 5’-CACCGTACTTTGCTGAATAAAGCTC-3’. The recombinant plasmid (14.5 μg), together with the packaging plasmids psPAX2 (14.5 μg) and pMD2.G (10 μg), were transfected into 70% confluence of HEK 293T cells in a 10 cm dish with Fugene (m/v=1:3) to package lentiviruses. The supernatants containing lentiviruses were collected at 48 h post-transfection and concentrated by centrifugation (2000 × rpm, 15 min). H1299 cells were then infected with lentiviruses containing 8 μg/ml polybrene. After 48 h.p.i., puromycin (2 μg/ml) was applied to select for G3BP knockout cells. The G3BP1 and G3BP2 stably knockout cells were obtained after 5-6 passages and the absence of G3BP1/2 expression was confirmed by Western blot analysis and genome sequencing.

### Construction of recombinant virus rIBV-nsp15-H238A

Plasmids pKTO-IBV-A, pGEM-IBV-B, pXL-IBV-C, pGEM-IBV-D, pGEM-IBV-E bearing IBV Beaudette fragment A, B, C, D and E covering the full-length genome (NC_001451.1) (see table 3) and plasmid pKTO-IBV-N containing N gene and 3’-UTR are a generous gifts from Prof. Shouguo Fang, Yangtze University. Nsp15-H238A mutation was introduced by using Mut Express II Fast Mutagenesis Kit V2 on pGEM-IBV-D (primers sequences were shown in Table 2). The *Bsa I/BsmB I* digested products of pKTO-IBV-A and pGEM-IBV-B were ligated by T4 ligase overnight, and the *Bsa I/BsmB I* digested products of C, D and E were ligated overnight. The AB and CDE were then ligated overnight to get the full-length cDNA genome with nsp15-H238A(AB+CDE). The full-length cDNA and *EcoRI* digested pKTO-IBV-N were subjected to *in vitro* transcription using T7 transcription kit (Promega), respectively, and added with cap structure using m7G (5’) ppp (5’) G RNA cap (New England biolabs). Next, the capped full-length RNA and IBV-N transcripts dissolved in 400 μl PBS were co-transfected into Vero cells by electroporation (450 v, 50 μF, 3 mSec, GenePulser Xcell, BIO-RAD). After 48 h, the supernatant was collected and used to inoculate new Vero cells. When syncytia appeared, the supernatant was collected again and passaged on Vero cells for 3 to 5 times. Finally, the virus-containing medium was collected and sequenced. Viral titer was determined by TCID_50_.

**Table 3.** Plasmids for rIBV nsp15-H238A construction

| Table 3 Plasmids for rIBV nsp15-H238A construction |  |  |  |  |  |
| --- | --- | --- | --- | --- | --- |
| Number | Plasmid | Resistance | Fragment | Size | Location (nt) |
| 1 | pKTO-IBV-A | Amp<sup>+</sup> | A | 6.4kb | 1-5751 |
| 2 | pGEM-IBV-B | Amp<sup>+</sup> | B | 3.0kb | 5752-8693 |
| 3 | pXL-IBV-C | Kan<sup>+</sup> | C | 6.8kb | 8689-15520 |
| 4 | pGEM -IBV-D | Amp<sup>+</sup> | D | 5.4kb | 15521-20900 |
| 5 | pGEM -IBV-E | Amp<sup>+</sup> | E | 6.7kb | 20887-27608 |

### Viral titer determination

Virus yield in the supernatant of rIBV-nsp15-H238A infected H1299 cells and G3BP1/2KO H1299 cells were determined by TCID_50_ assay. Briefly, the supernatant was serially diluted in 10-fold and inoculated 70% confluence of H1299 cells in 96 well plates. The cytopathogenic effect was observed after 4 days infection and the TCID_50_ was calculated by Reed-Muench method.

### Statistical analysis

The statistical analysis was analyzed with Graphpad Prism8 software. The data show as means ± standard deviation (SD) of three independent experiments. Significance was determined with Student’s test. *P* values < 0.05 were treated as statistically significant.

## Data availability statement

All relevant data are within the paper and its Supporting Information files.

## Acknowledgments

We acknowledge Prof Dingxiang Liu (South China University, China) for supplying the IBV Beaudette strain and the rabbit anti-IBV-N polyclonal antibody, and for his excellent scientific advice.

## Author Contributions

**Conceptualization:** Ying Liao, Bo Gao, Xiaoqian Gong, Maria Forlenza, Yingjie Sun

**Data Curation:** Ying Liao, Bo Gao, Lei Tan, Weiwei Liu

**Formal analysis:** Ying Liao, Bo Gao, Lei Tan

**Funding acquisition:** Ying Liao, Chan Ding

**Investigation:** Bo Gao, Xiaoqian Gong, Wenlian Weng

**Methodology:** Shouguo Fang

**Project Administration:** Ying Liao, Chunchun Meng, Xusheng Qiu

**Resource:** Shouguo Fang, Chunchun Meng

**Software:** Cuiping Song, Weiwei Liu

**Validation:** Ying Liao, Yingjie Sun, Chunchun Meng

**Visualization:** Yingjie Sun, Xusheng Qiu, Cuiping Song

**Supervision:** Ying Liao, Chan Ding, Maria Forlenza, Yingjie Sun

**Writing-original draft:** Bo Gao, Ying Liao

**Writing-review & editing:** Ying Liao, Maria Forlenza

## Correspondence address

Address correspondence and reprint requests to Prof Ying Liao, Waterfowl virus infectious diseases, Shanghai Veterinary Research Institute, Chinese Academy of Agricultural Sciences, Shanghai, 200241. P. R. China

## Abbreviations

Abbreviations used in this article:

IBV: infectious bronchitis virus
SG: stress granule
TLRs: Toll like receptors
RLRs: RIG-I like receptors
IFN: interferon
IRF3/7: interferon regulatory factor 3/7
PKR: double-stranded RNA-dependent protein kinase R
PERK: PKR-like endoplasmic reticulum kinase
GCN2: general control nonderepressible protein 2
HRI: heme-regulated inhibitor kinase
G3BP1/2: Ras GTPase-activating protein-binding protein 1/2
TIA-1: T cell-restricted intracellular antigen 1
TIAR: TIA-1-related protein
PABP1: poly(A) binding protein 1
RTC: replication and transcription complex
MEX3C: RNA-binding E3 ubiquitin protein ligase
Riplet: E3 ubiquitin protein ligase RNF135
DHX36: DEAH box protein 36
NEMO: NF-kappa-B essential modulator
ARS: sodium arsenite
MOI: multiplicity of infection
RT-PCR: reverse transcription-PCR
siRNA: small interfering RNA
h.p.i: hours post infection
ISG: interferon stimulated gene
IFIT1: interferon induced protein with tetratricopeptide repeats 1.

